# Combining multiple data sources in species distribution models while accounting for spatial dependence and overfitting with combined penalised likelihood maximisation

**DOI:** 10.1101/615583

**Authors:** Ian W. Renner, Julie Louvrier, Olivier Gimenez

## Abstract

1. The increase in availability of species data sets means that approaches to species distribution modelling that incorporate multiple data sets are in greater demand. Recent methodological developments in this area have led to combined likelihood approaches, in which a log-likelihood comprised of the sum of the log-likelihood components of each data source is maximised. Often, these approaches make use of at least one presence-only data set and use the log-likelihood of an inhomogeneous Poisson point process model in the combined likelihood construction. While these advancements have been shown to improve predictive performance, they do not currently address challenges in presence-only modelling such as checking and correcting for violations of the independence assumption of a Poisson point process model or more general challenges in species distribution modelling such as overfitting.
2. In this paper, we present an extension of the combined likelihood framework which accommodates alternative presence-only likelihoods in the presence of spatial dependence as well as lasso-type penalties to account for potential overfitting. We compare the proposed combined penalised likelihood approach to the standard combined likelihood approach via simulation and apply the method to modelling the distribution of the Eurasian lynx in the Jura Mountains in eastern France.
3. The simulations show that the proposed combined penalised likelihood approach has better predictive performance than the standard approach when spatial dependence is present in the data. The lynx analysis shows that the predicted maps vary significantly between the model fitted with the proposed combined penalised approach accounting for spatial dependence and the model fitted with the standard combined likelihood.
4. This work highlights the benefits of careful consideration of the presence-only components of the combined likelihood formulation, and allows greater flexibility and ability to accommodate real datasets.

## 1 Introduction

Species distribution models (SDMs), in which the distributions of species are modelled as a function of environmental predictors, rely on information about where a species has been observed (Guisan *et al.*, 2017). Different SDM methods have been developed over the past few decades to accommodate the different protocols by which this species information is collected. For example, logistic regression and its extensions are often used when species detections and non-detections are recorded at a set of systematically designed locations (known as “presence-absence” data), while point process models (PPMs, see Renner *et al*. (2015) for an overview) have emerged as a unifying framework for fitting SDMs informed by “presence-only” data, in which only information about species presence locations are available. Statistically, these methods are often fitted by maximising a corresponding likelihood expression, and the parameter estimates which maximise the likelihood may be used to produce maps of relative habitat suitability, reported as a habitat suitability index (Hirzel *et al.*, 2002), probability of species presence (Phillips *et al.*, 2006), or intensity of locations per unit area (Warton & Shepherd, 2010) depending on the method.

Increasingly, species data are available from multiple sources and types. Many papers have advocated for fitting models to a combination of the available data types, illustrating benefits in model performance (Miller *et al.*, 2019). Dorazio (2014) illustrated via simulations that adding a small amount of systematically-collected presence-absence data to available presence-only data significantly improves predictive performance. Fithian *et al*. (2015) showed that fitting a combined presence-only and presence-absence model to multiple species leverages the information of more abundant species to improve predictive performance for less prevalent species and allows sampling bias inherent in presence-only data to be estimated and corrected. These models are fitted by maximising a combined log-likelihood expression which is the sum of the log-likelihoods of the presence-only and presence-absence components:

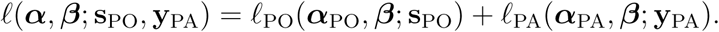

Here, **s**_PO_ contains the locations of a presence-only data source, while **y**_PA_ contains a vector of presence-absence detections and non-detections at a set of pre-selected sites. Parameters associated with the observation process unique to the presence-only and presence-absence data sets are denoted by ***α***_PO_ and ***α***_PA_, respectively, and collectively contained in the vector ***α***. Hereafter, we refer to these parameters as sampling bias parameters, as they may bias the intensity estimates as a result of the process of sampling the data. The key advancement of the combined likelihood approach is that the environmental response, parameterised by ***β***, is informed by both the presence-only and presence-absence data.

Such an approach implicitly assumes that the data sets are statistically independent, which allows for the combined log-likelihood to be expressed as a sum of the single-source log-likelihoods.

Other combinations may be done in similar fashion. For example, Koshkina *et al*. (2017) considered a combination of presence-only data with site-occupancy data, and Pacifici *et al*. (2017) developed a multivariate conditional autoregressive model to account for spatial autocorrelation in occurrence and detection error.

While these papers clearly advance the practice of fitting SDMs in important ways, they do not address some common challenges that arise in real datasets. For example, they all consider an inhomogeneous Poisson point process model (IPPPM) for the presence-only data in the combination. In many real data sets, however, the implicit assumption that the point locations are independently distributed conditional on the environment is not met. Residual clustering or repulsion of the point locations not accounted for with an IPPPM due to the observation process, unconsidered environmental covariates, or biological factors would hence render the IPPPM inappropriate. One option to account for spatial dependence is to consider a log-Gaussian Cox Process, as Gelfand & Shirota (2018) do for a combination of presence-only and presence-absence data. Furthermore, none of the current literature in combined likelihood approaches includes ways to account for possible overfitting that results from including too many covariates in the model.

However, advances in SDM literature provide solutions to these common problems. Diagnostic tools such as the inhomogeneous *K* function (Baddeley & Turner, 2000) and its simulation envelope (Diggle, 2003) can be used to determine departures from the independence assumption, and a wide number of alternative PPMs which account for spatial dependence may be included in the likelihood combination instead. Furthermore, penalised regression techniques such as the lasso penalty (Tibshirani, 1996) and its extension the adaptive lasso (Zou, 2006) may be used as a way to perform variable selection. Lasso regularisation has been shown to boost predictive performance of SDMs and has been applied to IPPPMs (Renner & Warton, 2013) and occupancy models (Hutchinson *et al.*, 2015).

In this paper, we present a penalised combined likelihood model in a way that it is more suitable for real data sets. In particular, we accommodate alternative forms of presence-only models to account for spatial dependence and affix a penalty on model complexity to address overfitting. In Section 2, we present the penalised combined likelihood formulation. In Section 3, we illustrate via simulations the improvements that this formulation provides and apply the proposed formulation to analyse the distribution of the Eurasian lynx (*lynx lynx*) in the Jura Mountains in eastern France. Finally, we present a discussion and further avenues for research in this area in Section 4.

## 2 Materials and Methods

### 2.1 Combined Penalised Likelihood Formulation

We define the weighted, combined penalised log-likelihood as follows

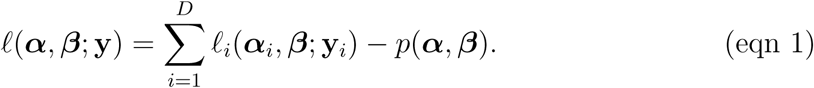

Here, ***α*** = (***α***_1_, …, ***α***_*D*_)^T^ is a *q*-dimensional vector that collects coefficients for the variables **Z** used to model sampling bias for each of the *D* components individually. The environmental response is measured by a set of variables **X** and is parametrised by ***β*** = (*β*_1_, …, *β*_*p*_)^T^, which is collectively informed by all *D* components. The species data for all *D* components is collected in a set **y**, with each individual data source **y**_*i*_ determining the form of the component likelihood ℓ_*i*_(***α***_*i*_, ***β***; **y**_*i*_). Finally, *p*(***α, β***) is a penalty term described in further detail below.

While many possibilities for the likelihood terms ℓ_*i*_(***α***_*i*_, ***β***; **y**_*i*_) are possible, we will focus on likelihood expressions for a PPM and for an occupancy model. For an IPPPM, we typically model the intensity of points *µ*(*s*) over a given study region *A* as a log-linear function of environmental variables **X** and sampling bias terms **Z** and derive estimates 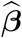 and 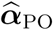 of the associated parameters by maximising a log-likelihood expression given by (Cressie, 1992):

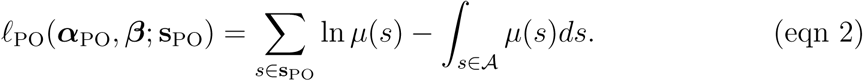

In the simple occupancy model we consider, each site *i* is visited *J*_*i*_ times. We collect the history of detections and non-detections for all *N* sites in a matrix **y**_occ_. We assume that the probability that site *i* is occupied is given by *ψ*_*i*_ and that the occupancy of the sites remains constant throughout the history of visits. We further assume the probability of detecting the species if present is *p*_*i*_. Under these assumptions, we can then model the probability of observing *y*_*i*_ detections at site *i* as

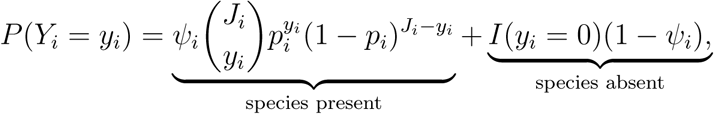

where *I*(·) is the indicator function.

We can relate the occupancy *ψ*_*i*_ of site *i* to an inhomogeneous Poisson intensity *µ*_*i*_ of the species distribution over site *i* as in Koshkina *et al*. (2017):

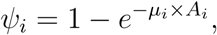

where *A*_*i*_ is the area of site *i*. Note that *µ*_*i*_ × *A*_*i*_ is an approximation of *∫*_*s*∈site *i*_ *µ*(*s*)*ds* that is reasonable if *µ*_*i*_ reasonably approximates the average intensity within site *i*.

As with the IPPPM, we can then model intensity as a log-linear function of environmental variables **X** and model detection probability *p*_*i*_ as a function of some detection covariates **Z**, such as the logit or complementary log-log function. We can then compute estimates 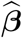 and 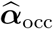 of the associated model parameters by maximising the log-likelihood expression given by:

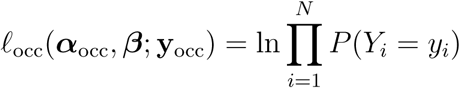

The term *p*(***α, β***) in eqn 1 is a penalty on model complexity applied to both the environmental parameters ***β*** and the sampling bias parameters ***α*** to shrink these parameters toward zero in order to boost predictive performance. Here, we consider both the traditional lasso penalty (Tibshirani, 1996) and the adaptive lasso penalty (Zou, 2006). For the traditional lasso penalty,

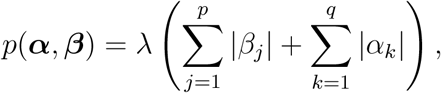

where *λ* is the tuning parameter. For the adaptive lasso penalty,

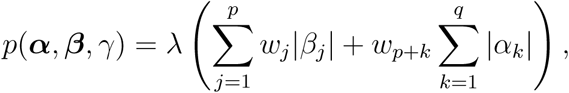

where **w** = (*w*_1_, …, *w*_*p*+*q*_)^T^ are weights for the adaptive lasso, typically of the form:

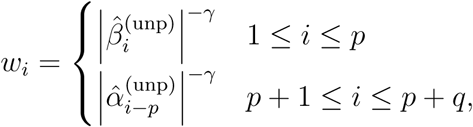

for *γ* > 0. Here, 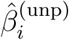 is the unpenalised coefficient estimate corresponding to the *i*^th^ environmental variable **x**_*i*_ and 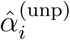 is the unpenalised coefficient estimate corresponding to the *i*^th^ sampling bias variable **z**_*i*_. The shape of the weights is determined by the parameter *γ*. The data-driven choice of the adaptive weights **w** ensures that more important covariates (*i*.*e*. those with coefficient estimates further away from 0) will be penalised less. This construction also enables the adaptive lasso to achieve so-called oracle properties (Zou, 2006), which means that asymptotically, the correct subset of coefficients will be chosen and the procedure has an optimal estimation rate.

We can use eqn 1 to represent the simpler framework introduced by Dorazio (2014) and Fithian *et al*. (2015) by setting *p*(***α, β***) = 0. We further extend this framework by considering alternative choices for those component likelihoods ℓ_*i*_(***α***_*i*_, ***β***; **y**_*i*_) informed by presence-only data. Rather than consider only inhomogeneous Poisson point process models, we consider area-interaction models (Widom & Rowlinson, 1970; Baddeley & van Lieshout, 1995) when diagnostic analysis of these data sources identifies spatial dependence among the presence-only locations. Area-interaction models account for spatial dependence through a vector of computed point interactions **t**_**s**_, which measure the proportion of overlap among circles of a nominal radius around the observed points **s**. They can account for both clustering and repulsion of points – the model parameter *η* characterises the nature of the spatial dependence, with values of *η* less than 1 signalling point repulsion and values of *η* greater than 1 signalling point clustering.

Because the likelihood expression of an area-interaction model is intractable, it is typically fitted via maximum pseudolikelihood (Besag, 1977):

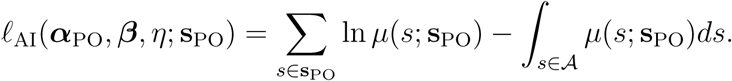

This log-pseudolikelihood expression appears the same as eqn 2, with the exception that the intensity *µ*(*s*) is replaced by conditional intensity *µ*(*s*; **s**_PO_) (Papangelou, 1974), reflecting the fact that for the area-interaction model, intensity at a location *s* is conditional on the other points in the pattern **s**_PO_.

### 2.2 Implementation in R

To fit models with the combined penalised log-likelihood in eqn 1, we have developed a set of functions in R inspired by the `optim` function and `ppmlasso` package (Renner & Warton, 2013). The main function `comb_lasso` takes as an input a list of species data, associated environmental data, and formulae for the environmental trend and sampling bias trends for each component, along with details such as type of presence-only likelihoods to use, the type of penalty, the number of models to fit, and the tuning parameter criterion. The function applies the coordinate descent algorithm of Osborne *et al*. (2000). This requires the derivatives of the component likelihoods (also known as “score equations”) to be computed. Analytical score equations are supplied directly to the `optim` function, which serves as the machinery of the optimisation. A tutorial illustrating use of this code for the simulations as performed in Section 3.1 as well as some functions written to plot intensity maps and features of the lasso penalisation is provided in the supplementary material.

## 3 Results

### 3.1 Simulations

To investigate the benefits of the proposed penalised combined likelihood formulation, we used the `rpoispp` function in `spatstat` (Baddeley & Turner, 2005) to generate a large inhomogeneous Poisson pattern **s**_true_ of roughly 10,000 points on a 30 × 30-unit square window from an intensity pattern defined by linear and quadratic terms of two generated variables (hence four meaningful covariates **x**_1_, …, **x**_4_ parameterised by coefficients *β*_1_, …, *β*_4_).

From this pattern, we generated two presence-only subsamples **s**_1_ and **s**_2_ biased by a different observation process. The first presence-only subsample **s**_1_ was biased by **z**_1_, the distance to a simulated road network, and the other **s**_2_ by **z**_2_, the distance to a simulated categorical covariate. We varied the size of the subsamples such that each pattern had 25, 100, or 400 points. We also varied the strength of the clustering of the presence-only subsamples by setting the coefficient of the interaction term *ν*_*i*_ = ln *η*_*i*_ for *i* = 1, 2. Here, the patterns either exhibit no clustering (*ν*_*i*_ = 0), moderate clustering (*ν*_*i*_ = 0.5) or strong clustering (*ν*_*i*_ = 1). In each case, the radius of interactions is set to 1 spatial unit. To sample the points in **s**_1_, we proceed as follows:

1. Initialise the set of sampled points **s**_1_ = ∅ and the point interactions 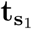 to be a vector of 0s
2. Compute the biased conditional intensity *µ*_1_(*s*; **s**_1_) at every point in **s**_true_ using **x**_1_, …, **x**_4_, the sampling bias covariate **z**_1_, and the current vector of point interactions 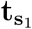, where the biased conditional intensity is defined as follows:

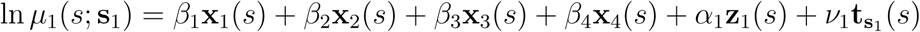
3. Set *µ*_1_(*s*; **s**_1_) = 0 for all *s* ∈ **s**_1_. That is, we set the conditional intensity for any point already selected in **s**_1_ to 0 to ensure these points are not resampled
4. Randomly select a point from **s**_true_ with sampling probabilities proportional to the conditional intensities and add the selected point to **s**_1_
5. Update the vector of point interactions 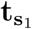 for all points in **s**_true_ using the internal `evalInteraction` function in `spatstat`, which computes point interactions based on a supplied point pattern for a given set of locations and interaction radius
6. Repeat steps 2-5 until we have sampled the desired number of points

We sample **s**_2_ in a similar manner, using **z**_2_ instead of **z**_1_ to create the sampling bias and computing point interactions 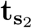.

Because the true pattern **s**_true_ is Poisson, this simulation setup emulates a scenario in which the clustering of the observed point patterns is an artefact of the observation process – this can happen if, for example, records are publicly available and enthusiasts for the species report further observations near the publicly available locations (Johnston *et al.*, 2019).

We also generated a history **y**_occ_ of detections and non-detections from 5 visits to each of 100 sites centred along a regular grid in the 30 × 30-unit observation window to emulate a data set for which we could consider occupancy modelling. The species was considered present at a site if the closest point in the pattern **s**_true_ was within a distance of 0.18 units of the centre of the site, such that the area of each site is roughly 0.1 square units. The history of detections and non-detections at each site where the species was considered present was randomly generated according to detection probabilities defined by the inverse of the cloglog function evaluated at a generated detection covariate **z**_3_.

Finally, we generated four dummy covariates **d**_1_, …, **d**_4_ to include in fitted models that were meaningless in describing the true species distribution. We did this to reflect the fact that in real applications, we may not know which among a suite of candidate variables truly determine the species distribution. We ensured that the maximum absolute correlation among all pairs of variables was smaller than 0.5.

After generating the species data, we fit a number of models, using as input environmental covariates the four meaningful covariates **x**_1_, …, **x**_4_ (parameterised by *β*_1_, …, *β*_4_) as well as four dummy covariates **d**_1_, …, **d**_4_ (parameterised by *β*_5_, …, *β*_8_) and using as sampling bias covariates **z**_1_, **z**_2_, and **z**_3_ (parameterised by *α*_1_, *α*_2_, and *α*_3_). For both Poisson and area-interaction presence-only likelihoods, we fit a model without any penalty, with a lasso penalty, and with an adaptive lasso penalty. For the models fitted with either a lasso or an adaptive lasso penalty, we fit regularisation paths of 1000 models, increasing the penalty from 0 to the smallest penalty *λ*_max_ that would shrink all coefficients to 0, thus covering the entire scope of possible model sizes. The model which minimised BIC was chosen among the 1000 fitted models. We considered as species data a combination of all three of **s**_1_, **s**_2_, and **y**_occ_. This led to a total of six models being fitted, summarised in Table 1.

**Table 1:** Models fitted in each simulation using the proposed combined penalised likelihood. The models also varied based on the likelihood expression for any presence-only components and the type of penalty used, if any.

| Model | Species Data | Presence-only likelihood | Penalty |
| --- | --- | --- | --- |
| 1 | $\mathbf{s}_1, \mathbf{s}_2$, and $\mathbf{y}_{\text{occ}}$ | IPPPM | None |
| 2 | $\mathbf{s}_1, \mathbf{s}_2$, and $\mathbf{y}_{\text{occ}}$ | IPPPM | Lasso |
| 3 | $\mathbf{s}_1, \mathbf{s}_2$, and $\mathbf{y}_{\text{occ}}$ | IPPPM | Adaptive Lasso |
| 4 | $\mathbf{s}_1, \mathbf{s}_2$, and $\mathbf{y}_{\text{occ}}$ | Area-interaction | None |
| 5 | $\mathbf{s}_1, \mathbf{s}_2$, and $\mathbf{y}_{\text{occ}}$ | Area-interaction | Lasso |
| 6 | $\mathbf{s}_1, \mathbf{s}_2$, and $\mathbf{y}_{\text{occ}}$ | Area-interaction | Adaptive Lasso |

To evaluate performance, we compared the integrated mean squared error of the true intensity surface with rescaled fitted intensity surfaces of the six models. The fitted intensity surfaces were rescaled to have the same mean intensity as the true intensity surface to ensure that fair comparisons are made as models using different species data sources will have varying intercepts to reflect the estimated abundance of the points.

We performed 1,000 simulations of the data sets for each of the nine combinations of presence-only data set size and clustering strength and the resultant model fits on 512GB nodes powered by 3.0 GHz Intel Xeon Gold (E5-6154) processor from the University of Newcastle’s High Performance Computing cluster. The 9,000 simulation tasks took approximately 7,000 hours.

Figure 1 shows boxplots of the calculated integrated mean squared errors from the simulations. From these results, we can draw the following conclusions. First, the models fitted with the area-interaction presence-only likelihoods have performance benefits over the models fitted with Poisson presence-only likelihoods when clustering is present. When clustering is not present (first column), a setting for which the Poisson likelihood is appropriate, the models fitted with area-interaction presence-only likelihoods perform no worse than models fitted with Poisson presence-only likelihoods. Comparing the plots across rows and down columns, we see that the performance advantage of the models fitted with area-interaction presence-only likelihoods tends to increase as the degree of clustering gets larger and as the sample size increases, respectively.

**Figure 1:**
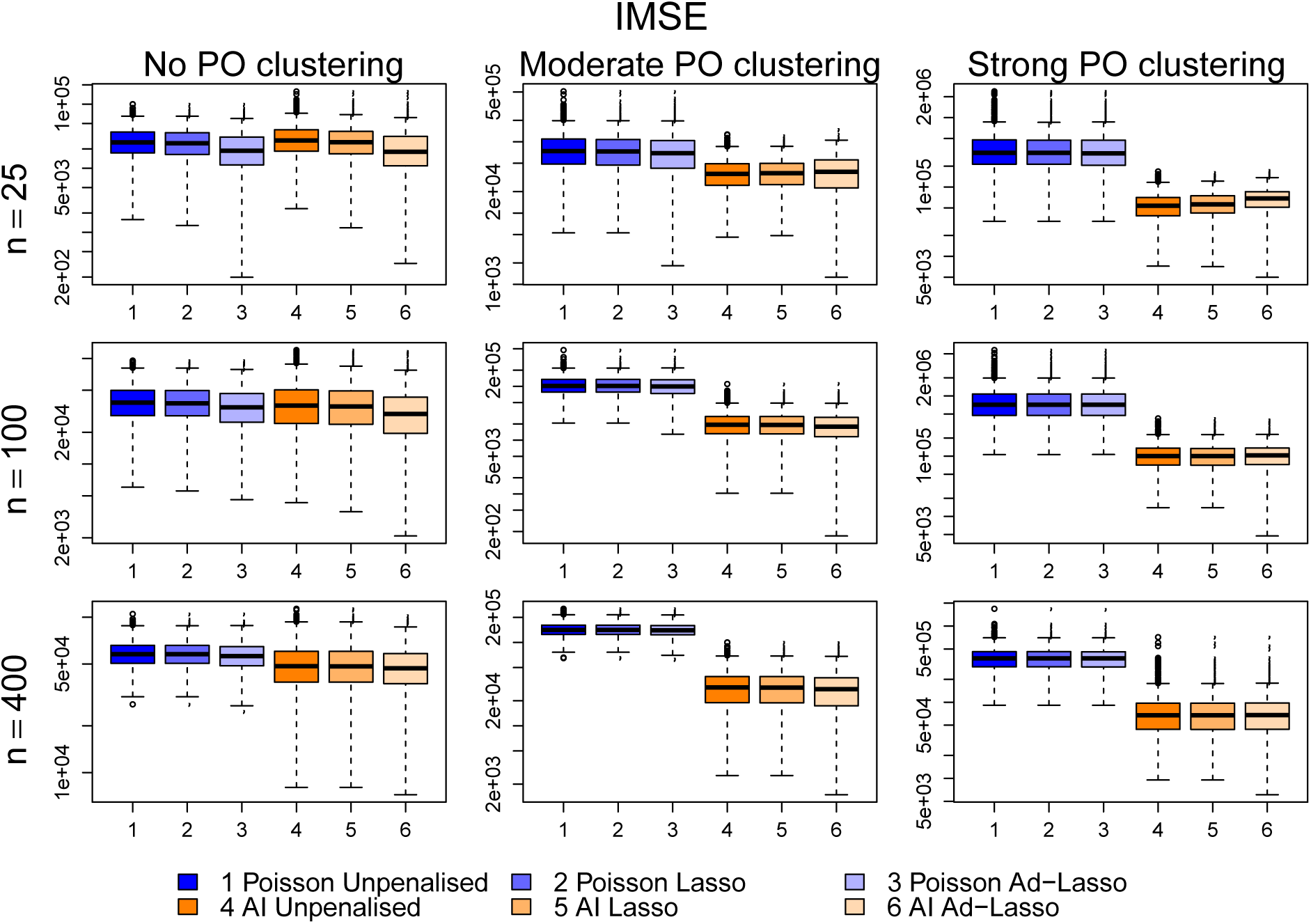
Boxpots of integrated mean squared error for the six models described in Table 1 for different combinations of presence-only sample size and clustering strength.

In the Appendix, we show that the parameter coefficients *β*_1_, …, *β*_4_ corresponding to the meaningful covariates **x**_1_, …, **x**_4_ are increasingly biased away from 0 for the models fitted with Poisson presence-only likelihoods, both as sample size increases and as the strength of presence-only clustering increases. The inclusion of the area-interaction term takes an increasing amount of signal from the environmental covariates as the strength of the presence-only clustering increases. For low sample sizes, there is a suggestion that this signal dampening may be too strong, though such an overcorrection disappears as sample size increases.

Second, penalisation via the lasso or adaptive lasso improves model performance when there is no presence-only clustering, and this improvement is greatest for smaller sample sizes. This is an expected conclusion given the danger of overfitting is greater with fewer observations. Models penalised with the adaptive lasso tend to outperform models penalised with the lasso when there is no presence-only clustering. However, lasso penalisation does not notably improve performance when there is presence-only clustering. In fact, there is a suggestion that applying a lasso penalty may slightly hinder performance when applying an area-interaction presence-only likelihood for small sample sizes. Although the benefits of penalisation are negligible with large data sets, fitting models with a penalty does not hurt the performance.

In summary, it appears that the proposed combined penalised likelihood framework provides the best performance. Furthermore, incorporating area-interaction presence-only likelihoods improves performance when clustering is present, and can likewise reliably estimate that there are negligible point interactions if clustering is not present, in effect relaxing to the simpler model with Poisson presence-only likeihoods when this additional complexity is not needed. A more detailed discussion of the simulation results, including boxplots of the fitted coefficients, appears in the Appendix.

### 3.2 Analysis of Eurasian lynx distribution in the Jura Mountains

We now demonstrate the use of the combined penalised likelihood approach to analyse the distribution of the Eurasian lynx in the Jura Mountains in eastern France.

Lynx went extinct in France at the end of the 19th century due to habitat degradation, human persecution and decrease in prey availability (Vandel & Stahl, 2005). The species was reintroduced in Switzerland in the 1970s (Breitenmoser *et al.*, 1998), then re-colonised France through the Jura mountains in the 1980s (Vandel & Stahl, 2005). The species is listed as endangered under the 2017 IUCN Red list and is of conservation concern in France due to habitat fragmentation, poaching and collisions with vehicles. The Jura holds the bulk of the French lynx population.

We have three sources of lynx data in the Jura Mountains: a presence-only data set consisting of 440 opportunistic sightings in the wild from 2009-2011 (denoted **s**_w_), another presence-only data set consisting of 240 reported interferences of lynx with domestic livestock in 2009-2011 (denoted **s**_d_), and pictures of lynx taken from cameras set up in 73 locations **s**_c_ in the Jura Mountains in 2012. Lynx presence-only data were made of presence signs sampled all year long thanks to a network of professional and non-professional observers. Every observer is trained during a 3-day teaching course led by the French National Game and Wildlife Agency (ONCFS) to document signs of the species’ presence (Duchamp *et al.*, 2012). Presence signs went through a standardised control process to prevent misidentification (Duchamp *et al.*, 2012). The camera data has daily reportings of the lynx across a total of 77 days. Due to this, we can consider the picture history of lynx at the camera locations in an occupancy modelling framework (Blanc *et al.*, 2014). In particular, we split the 77-day period into seven 11-day periods, such that the site history **y**_c_ comprises seven detections and non-detections at each site in **s**_c_ over each 11-day period.

Figure 2 shows the locations of the sightings in both presence-only data sets as well as the locations of the cameras. Both presence-only data sources appear to have different distributions, reflecting different sampling biases. There are more wild sightings in the northeast of the Jura Mountains, and more domestic interferences toward the southwest. Additionally, there appear to be some tight clusters within both data sets, with several records very close to each other.

**Figure 2:**
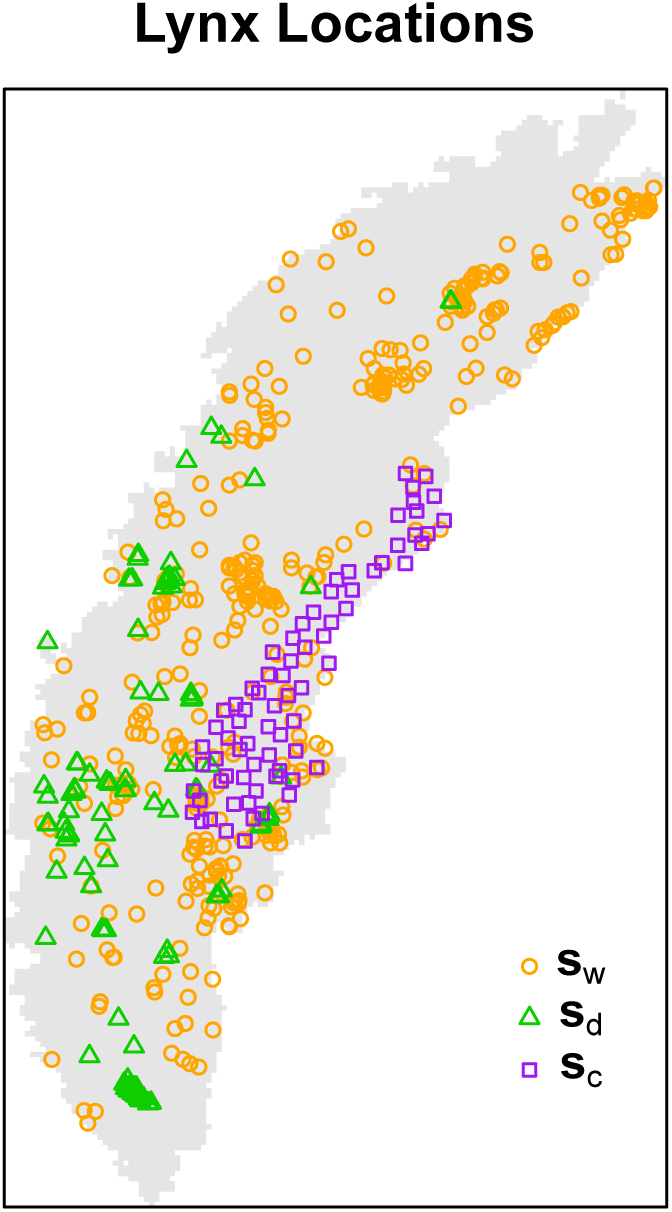
Locations of the lynx data in the Jura Mountains, including 440 observations in the wild **s**_w_, 220 reports of domestic interference **s**_d_, and 73 camera traps **s**_c_.

To model the lynx distribution, we consider altitude, percentage of forest cover, distance to the nearest water source, and human population density as environmental variables. We model sampling bias in the wild records **s**_w_ with distance to the nearest main road and distance to the nearest train line, and sampling bias in the domestic records **s**_d_ with distance to the nearest farm and percentage of agricultural land. Finally, we model detection probability for the camera data with distance to the nearest urban area. We established this set of potential candidate environmental and detection variables based on previously studied species habitat preferences and detectability (Bouyer *et al.*, 2015). The Corine Land Cover land use repository from 2012 (Büttner *et al.*, 2014) supplies a map of land coverage including urban areas, water areas, forest areas, farm areas, and agricultural areas that was used to generate the percentage of forest areas and agricultural areas over 1 km × 1 km cells as well as distances to the nearest urban area, water source, and farm. Altitude was averaged over 1km × 1km cells from data available in the `raster` package in R, while human population density was averaged over 1 km × 1km cells taken from version 4 of the Gridded Population of the World data repository (Center for International Earth Science Information Network (CIESIN) – Columbia University, 2016). Distances from the nearest main road and railway were computed from shapefiles from Version 151 of the ROUTE 500 database, accessible at http://professionnels.ign.fr/route500.

We fitted initial separate IPPPMs to the wild records **s**_w_ and the domestic records **s**_d_ using linear, quadratic, and interaction terms for the four environmental covariates, and linear terms for the sampling bias covariates. From these models, we are able to assess whether the assumption of independence inherent to the IPPPMs is appropriate with simluation envelopes of the inhomogeneous *K*-function in `spatstat`, as shown in Figure 3. Both of the envelopes for the IPPPMs fitted to the wild model (left panel) and the domestic model (middle panel) demonstrate additional clustering as the observed inhomogeneous *K*-function values plotted in red fall above the simulation envelopes for small radii. This suggests that fitting an IPPPM is inappropriate for these data sets. The right panel shows a simulation envelope of the cross *K* function as produced by the `Kcross.inhom` function of `spatstat`, which counts the expected number of wild sightings within a given distance of a domestic sighting, conditional on the spatially varying intensities of both patterns. We estimate the wild and domestic intensities from area-interaction models, and as the observed values of the cross *K*-function fall within the envelope boundaries, this suggests that there is no clustering across the two data sets. This, in turn, suggests that the observed clustering within the wild and domestic data sets may be more likely attributable to the observation process than to some biological reality that induces clustering or a missed environmental covariate.

**Figure 3:**
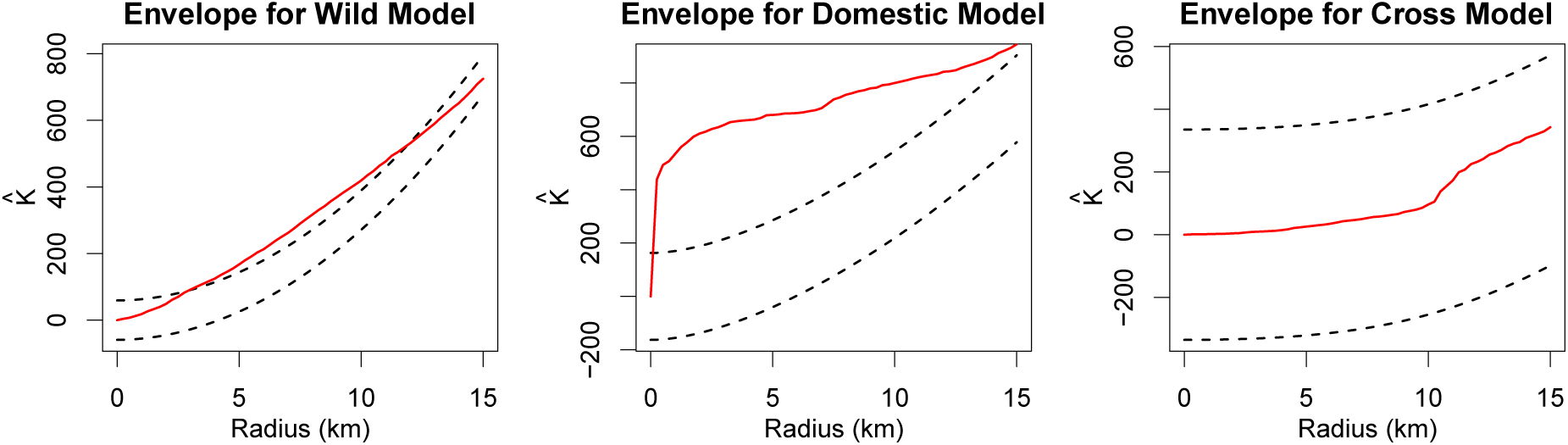
95% simulation envelopes of the inhomogeneous *K*-function for the fitted IPPPM of the wild records (left), the fitted IPPPM of the domestic records (middle), and across the two fitted IPPPMs (right).

Consequently, we fit combined likelihood models using both the standard, unpenalised approach (analogous to Model 1 in Table 1) and the combined penalised likelihood formulation eqn 1 with a lasso penalty and area-interaction models for the presence-only data sources (analogous to Model 5 in Table 1). The radii chosen to capture the residual spatial patterning in the wild and domestic models are 2km and 5km, as chosen by the `profilepl` function in `spatstat`.

Figure 4 shows the bias-corrected fitted intensities from these two models. For the combined model which uses IPPPMs (left panel), the fitted intensity is corrected for the sampling bias terms modelled for the presence-only components using the method of Warton *et al*. (2013). For the combined penalised model which uses area-interaction models (right panel), the fitted intensity is corrected for these same sampling bias terms as well as the fitted point interactions – that is, we treat the interaction parameter *ν* as belonging to the set of sampling bias parameters ***α***. The fitted models show strikingly different patterns, with the model which uses area-interaction components highlighting much more of the Jura Mountains as preferred habitat of lynx than the model which uses IPPPMs. The models suggest similar numbers of points throughout the Jura, but the distribution of these points are more heavily concentrated in the IPPPM model. This is because the area-interaction terms in the AI model lessen the impact of some clusters of points on the scale of the displayed bias-corrected intensities.

**Figure 4:**
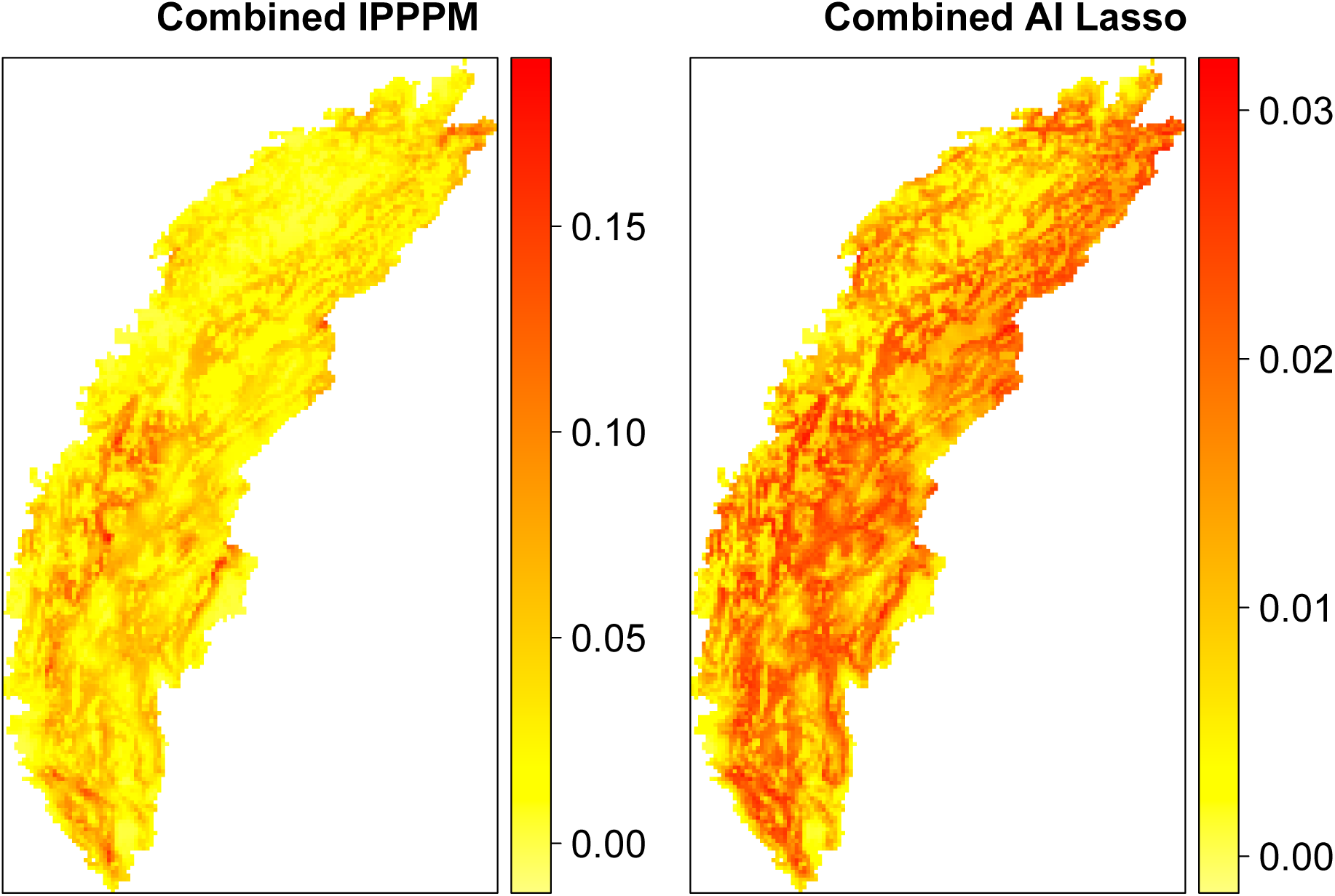
Fitted intensities using the combined likelihood formulation. Left: the model is fitted without any penalty and using inhomogeneous Poisson point process models for the presence-only data sources. Right: the model is fitted with a lasso penalty and using area-interaction models for the presence-only sources.

We do not have access to additional data with which to validate the performance of these models such as GPS data as in Gould *et al*. (2019), but the results of Section 3.1 suggest that the model which uses area-interaction components is likely to better reflect the true distribution of lynx.

The combined penalised model with the area-interaction components found the optimal lasso penalty was 0, resulting in a model which included all 18 covariates and both of the area-interaction terms. The fact that the optimal penalty is 0 suggests that the suite of covariates we chose to include, motivated by existing literature, seems to have been a good choice. In general, we recommend use of the lasso penalty as a safeguard against overfitting, particularly in contexts where the suite of candidate covariates for a species is less established as an insurance against overfitting.

## 4 Discussion

The proposed combined penalised likelihood framework addresses some common problems that arise in real datasets. The flexibility to incorporate an area-interaction likelihood when there is spatial dependence in the presence-only data set and affix a penalty on model complexity enables improvements in predictive performance, as shown in Section 3.1.

### 4.1 Possible extensions

Despite these improvements, further advances are possible. Other penalty structures could be incorporated into the same framework. While the lasso and adaptive lasso showcased here show clear benefits in simulations, other penalised likelihood variants such as SCAD (Fan & Li, 2001) could lead to superior performance in some situations, and alternative methods to BIC of choosing the size of the penalty such as the Extended Bayesian Information Criterion (“ERIC”, Hui *et al.*, 2015) could likewise be used.

While we make use of the area-interaction likelihood in this paper, there is a large family of Gibbs PPMs (Cressie, 1992) which accommodate different sorts of spatial dependence that could be used. Our choice of the area-interaction model as the alternative is motivated by the fact that it accommodates interactions of all orders instead of just pairwise interactions and that it can be used to model both clustering and repulsion of points.

The inclusion of the area-interaction terms dampens the signal of the environmental covariates. Although this makes sense when spatial dependence exists, we may dampen the signal too much. In the context of species distribution models, we might ask the question, “Does a given species record exist because its location is in particularly suitable habitat for the species, or because there are other records nearby?” If the answer to this question appears to be “both”, as is often the case for presence-only data, we are at risk of “spatial confounding”. In the single-source context, Hodges & Reich (2010) propose restricting the spatial effect to be orthogonal to the fixed covariate effects, while Simpson *et al*. (2017) and Sørbye *et al*. (2019) suggest careful selection of associated spatial priors to alleviate this risk. With our implementation, we could achieve something similar to the latter two papers by adjusting the magnitude of the lasso penalty for the area-interaction terms. In the Appendix, we highlight the tradeoff between the estimates of the interaction parameters 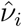 and both the estimates of the environmental parameters 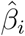 and the sampling bias parameters 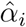. However, a full exploration of the effects of spatial confounding remains an open area of research and is beyond the scope of this paper.

In both the simulations in Section 3.1 and the lynx data analysis in Section 3.2, we made the rather limiting assumption of a closed population and that sites are either always occupied or always unoccupied. Nonetheless, occupancy models which take into account changing site dynamics could be used (MacKenzie *et al.*, 2003). Similarly, we have ignored the temporal aspect of the lynx distribution in this paper, but there is a wide suite of tools to fit spatio-temporal models in order to capture distribution dynamics for both the aforementioned occupancy modelling component as well as presence-only components (Cressie & Wikle, 2015).

Further improvements could be made by incorporating source weights in situations in which the data sources vary in quality. Indeed, presence-only data sources may be more prone to errors in coordinate locations as well as correct species identification, as they often include records by amateur enthusiasts. The combined penalised likelihood framework could easily be extended to include weights for the various data sources by adding a vector of source weights **w** = (*w*_1_, …, *w*_*D*_)^T^ to the formulation in eqn 1:

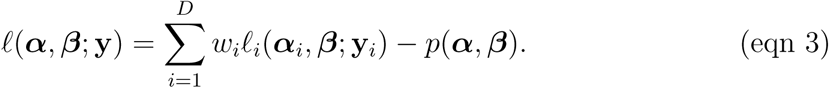

One possible strategy to incorporate such weights in eqn 3 could be to compare performance of single source models on independent data and upweight the contribution of data sources that are shown to have good performance.

Finally, while we incorporate sampling bias as a linear effect, non-linear effects can also be used as appropriate for a given sampling protocol, for example with distance sampling as discussed in Yuan *et al*. (2017).

### 4.2 Accounting for dependence within and among data sources

In the lynx data analysis in Section 3.2, we diagnosed spatial dependence within each of the presence-only data sources but found no spatial dependence across data sources. Tools such as the inhomogeneous *K*-envelope provide great insight into the underlying individual spatial processes that are observed. However, such diagnostic tools are not currently available for the combined likelihood models, and research in this area would be valuable as these models grow in popularity.

Another approach to constructing SDMs from multiple data sources could be to introduce a common latent spatial term *ξ*(*s*), such as a Gaussian random field, which would account for spatial dependence among points in all of the data sources as in Gelfand & Shirota (2018). The resulting likelihood expression would be:

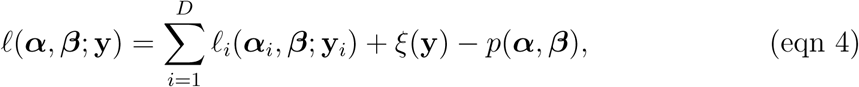

where *ξ*(**y**) ∼ MVN(**0, ∑**). Models of this type are typically fitted in a Bayesian framework. We could reduce the dimension of *ξ* through methods like fixed rank kriging or induce sparsity in **∑** through lasso-type penalties such that the likelihood in eqn 4 could be fitted with software such as Template Model Builder (TMB, Kristensen *et al.*, 2016). Another way to achieve sparsity is with the stochastic partial differential equation approach (SPDE, Lindgren *et al.*, 2011), as implemented in the `inlabru` package (Bachl *et al.*, 2019).

### 4.3 Conclusion and Perspectives

The development of statistical methods is often motivated by new challenges raised by novel types of data sets. While the current literature on combined likelihood approaches represents a significant recent advancement, advances in other areas can be lost if not carried over with such methodological developments. This paper attempts to build a bridge between this exciting new arena for species distribution modelling and the rich suite of tools available for species distribution modelling, particularly that for presence-only data. Our hope is that other such bridges continue to be built in this spirit.

## Supporting information

Optim Lasso demo

## 5 Acknowledgements

We thank the staff from the French National Game and Wildlife Agency, the Forest National Agency and the Departmental Federation of Hunters of Jura department, who collected the photographs during the camera-trapping session. We also thank all the volunteers that are members of the Réseau Loup-Lynx that collect every year precious presence signs of lynx all over its distribution area. We thank the University of Newcastle through the Early to Mid-Career Researcher Visiting Fellowship and the Université Paul-Valéry Montpellier through the Professeurs en mobilité universitaire fund for funding visits for I.R. which helped facilitate the collaborations that resulted in this work. O.G. was funded by CNRS and the “Mission pour l’Interdisciplinarité” through the “Osez l’Interdisciplinarité” initiative. Finally, we thank the reviewers for their very helpful comments, which have helped us greatly improve the paper.

## 6 Authors’ Contributions

I.R. and O.G. conceived the concept of the paper. I.R. developed the code to fit the models. J.L. sourced the species coordinates and covariates for the lynx analysis. I.R. and O.G. wrote the manuscript. I.R. and J.L. developed the tutorial in the supplementary information. All authors were involved in editing drafts of the manuscript.

