## Supplementary material for "Combining multiple data sources in species distribution models while accounting for spatial dependence and overfitting with combined penalised likelihood maximisation": Supp material

### Appendix: Further results and analysis of simulations

In Section 3.1 of the main manuscript, we simulated two presence-only data sets and one occupancy data set to mimic the setting of the lynx data set described in Section 3.2. The simulations suggested that the combined models that incorporated an area-interaction presence-only likelihood had better predictive performance than models that incorporated a Poisson presence-only likelihood, and that this performance benefit increased with the degree of presence-only clustering and sample size. Here, we explore these results further by examining parameter estimation.

Figure A1 contains boxplots for the four coefficients $\hat{\beta}_1, \ldots, \hat{\beta}_4$ corresponding to the four meaningful environmental covariates $\mathbf{x}_1, \ldots, \mathbf{x}_4$ for each of the six models fitted to the data. The true values used to generate the species patterns $\mathbf{s}_1$ and $\mathbf{s}_2$ were $\beta_1 = 0.5$, $\beta_2 = -0.3$, $\beta_3 = 0.5$, and $\beta_4 = -0.3$. By looking down each column of plots, we see that the estimates become increasingly biased away from 0 for the models which incorporate a Poisson presence-only likelihood. This makes sense when there is presence-only clustering, as the environmental covariates absorb signal from the clustering as well as the environmental signal. Interestingly, this overinflation occurs even when there is no presence-only clustering, though not to the same degree.

The inclusion of the area-interaction terms increasingly mitigates this overinflation as the strength of presence-only clustering increases. The combined models with area-interaction presence-only likelihoods initially appear to overcorrect when $n = 25$ such that the estimates are biased toward 0, but then undercorrect when $n = 400$. This suggests that additional data is tending to amplify the environmental signal, and indicates potential spatial confounding, as mentioned in the discussion of the main manuscript. Nonetheless, the inclusion of the area-interaction terms leads to less biased parameter estimates overall, particularly as the strength of presence-only clustering and sample size increases. This coincides nicely with the simulation results presented in Figure 1 of the main manuscript.

Figure A2 contains boxplots of the coefficient estimates $\hat{\beta}_5, \ldots, \hat{\beta}_8$ corresponding to the dummy variables $\mathbf{d}_1, \ldots, \mathbf{d}_4$. This plot suggests that the lasso models and adaptive lasso models (the lighter colored boxplots within both the models with Poisson and area-interaction presence-only likelihoods) are more concentrated around the true value of 0, as expected. Once again, the medians of the models fitted with Poisson presence-only likelihoods are being dragged away from 0 as sample size increases, though the variances are smaller. The estimates are increasingly concentrated around 0 as the strength of

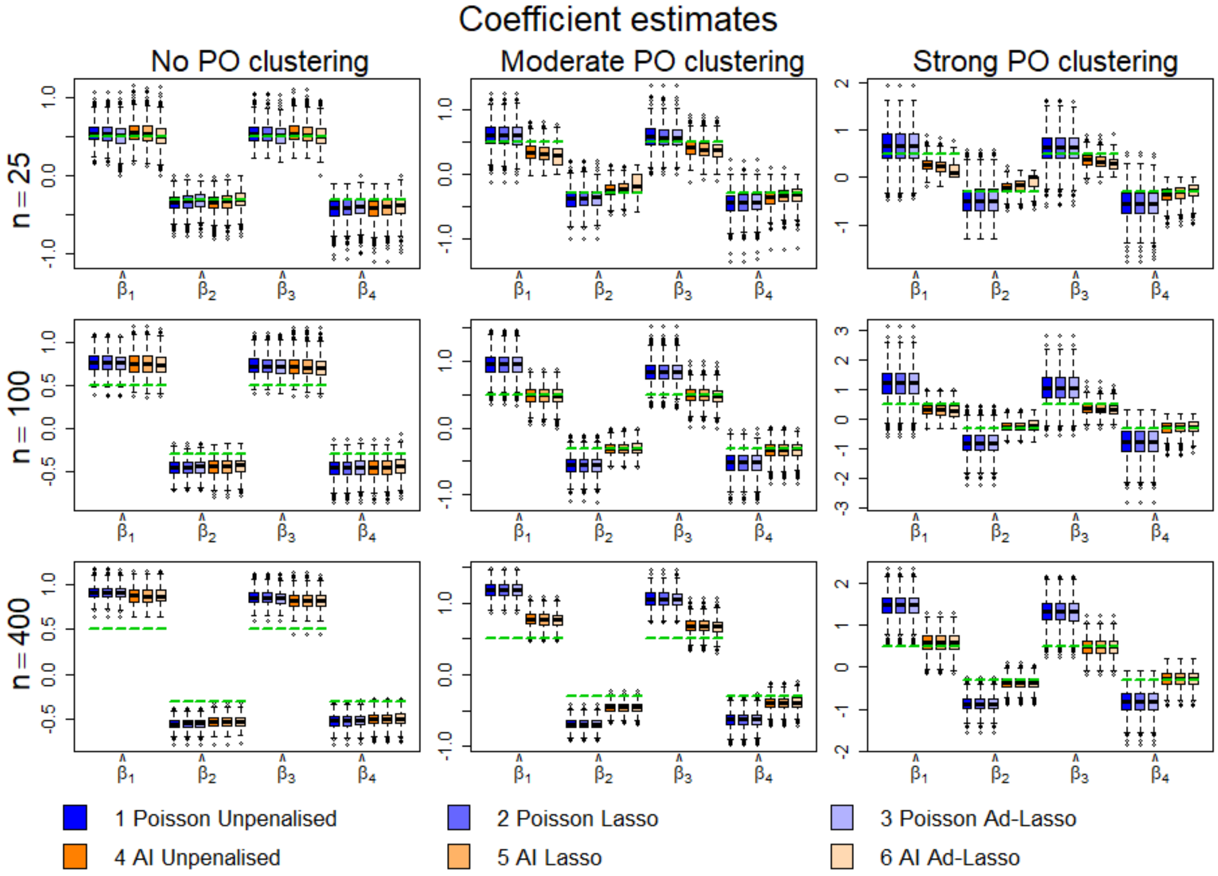


Figure A1: Boxpots of coefficient estimates corresponding to the four meaningful environmental covariates for the six models described in Table 1 of the manuscript. The green lines indicate the true values of the parameters used to generated the simulated data.

presence-only clustering increases for the models fitted with area-interaction presence-only likelihoods. This makes sense as the area-interaction terms dampen the signal of the other covariates.

Figure A3 contains boxplots of the coefficient estimates $\hat{\alpha}_1$ and $\hat{\alpha}_2$ corresponding to the two observer bias covariates $\mathbf{z}_1$ and $\mathbf{z}_2$ as well as the estimates $\hat{\nu}_1$ and $\hat{\nu}_2$ corresponding to the area-interaction terms. For the simulations, the true values of the observer bias covariates were $\alpha_1 = 1$ and $\alpha_2 = 1$. The true values of the area-interaction coefficients were $\nu = 0$ for the patterns with no presence-only clustering, $\nu = 0.5$ for the patterns with moderate presence-only clustering, and $\nu = 1$ for the patterns with strong presence-only clustering.

We see that the observer bias parameters $\alpha_1$ and $\alpha_2$ are better estimated with larger sample sizes, and also better estimated by the models that incorporate Poisson presence-only likelihoods, particularly as the strength of the presence-only clustering increases. This superior estimation is likely attributable to spatial confounding – the estimates in

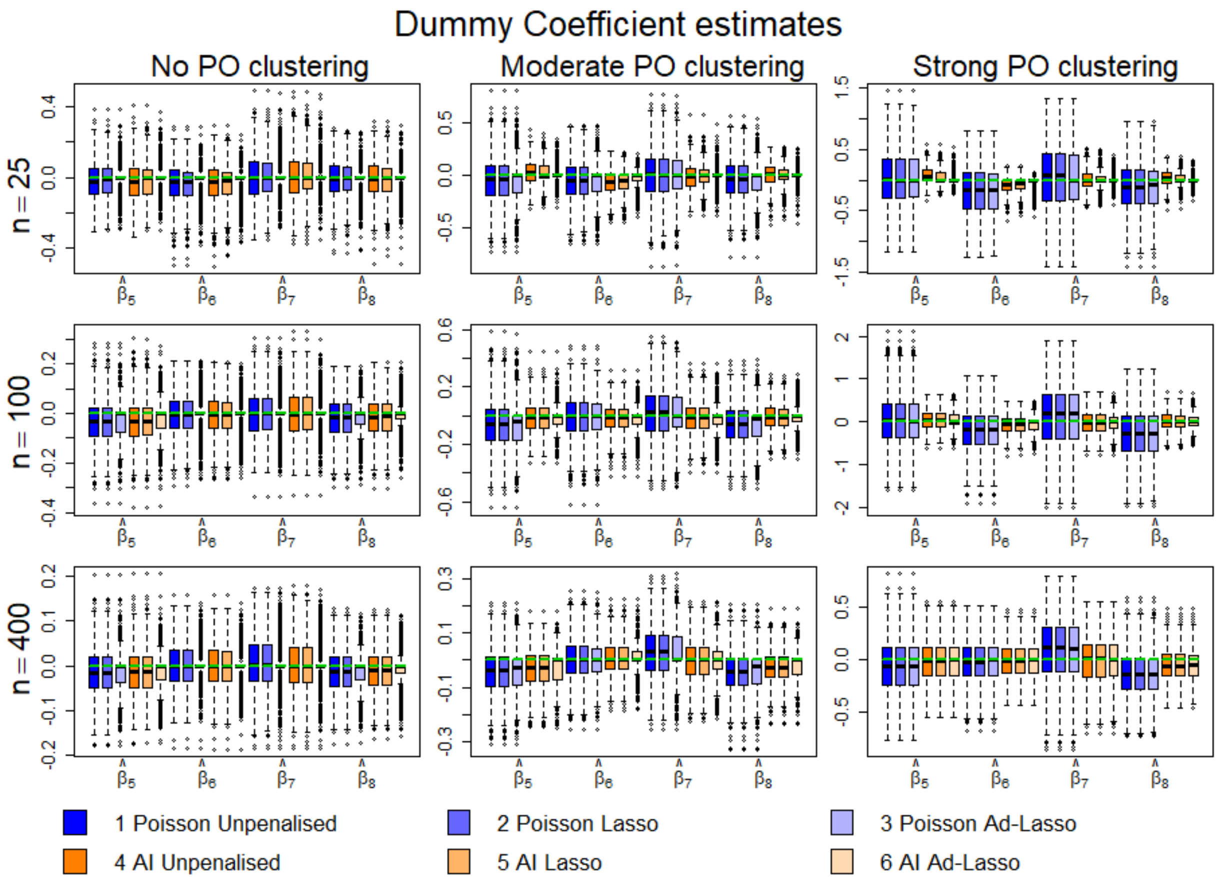


Figure A2: Boxpots of coefficient estimates corresponding to the four dummy environmental covariates for the six models described in Table 1 of the manuscript. The green lines indicate the true values of the parameters used to generated the simulated data.

the models fitted with Poisson presence-only likelihoods do not compete with estimates of the interaction terms for signal as they do in the case of the models fitted with area-interaction likelihoods.

The estimates of the interaction coefficients are naturally superior for the models fitted with area-interaction presence-only likelihoods. As sample size increases, the estimates of the interaction coefficients become larger, being too strong for $n = 100$ and $n = 400$. It seems there is a tradeoff between the estimation of the observer bias variables and the area interaction covariates. This makes sense – these variables will naturally be negatively correlated. In the case of the first presence-only data source, points are simulated to be both close to roads and close to each other. From a bias perspective, this means we can ask of a particular point, “was this point sampled because it was near to a road or because it is near to other points?” Similarly for the second presence-only data source, points are simulated to be both close to a particular level of a categorical covariate and close to each other. As Figure A4 indicates, these negative correlations grow stronger with increasing sample size, making it difficult for the models to disentangle the effects of the observer bias covariates and the area-interaction terms even with the additional data.

As both the area-interaction terms and the observer bias variables are treated in the same way when making predictions, it is the overall contribution of both the observer bias and the area-interaction terms that defines the estimated bias surface corrected by the method of Warton *et al.* (2013). With the high negative correlations, overestimation of the area-interaction contribution may lead to underestimation of the contribution of the observer bias variables, but the overall bias surface may still be reasonable.

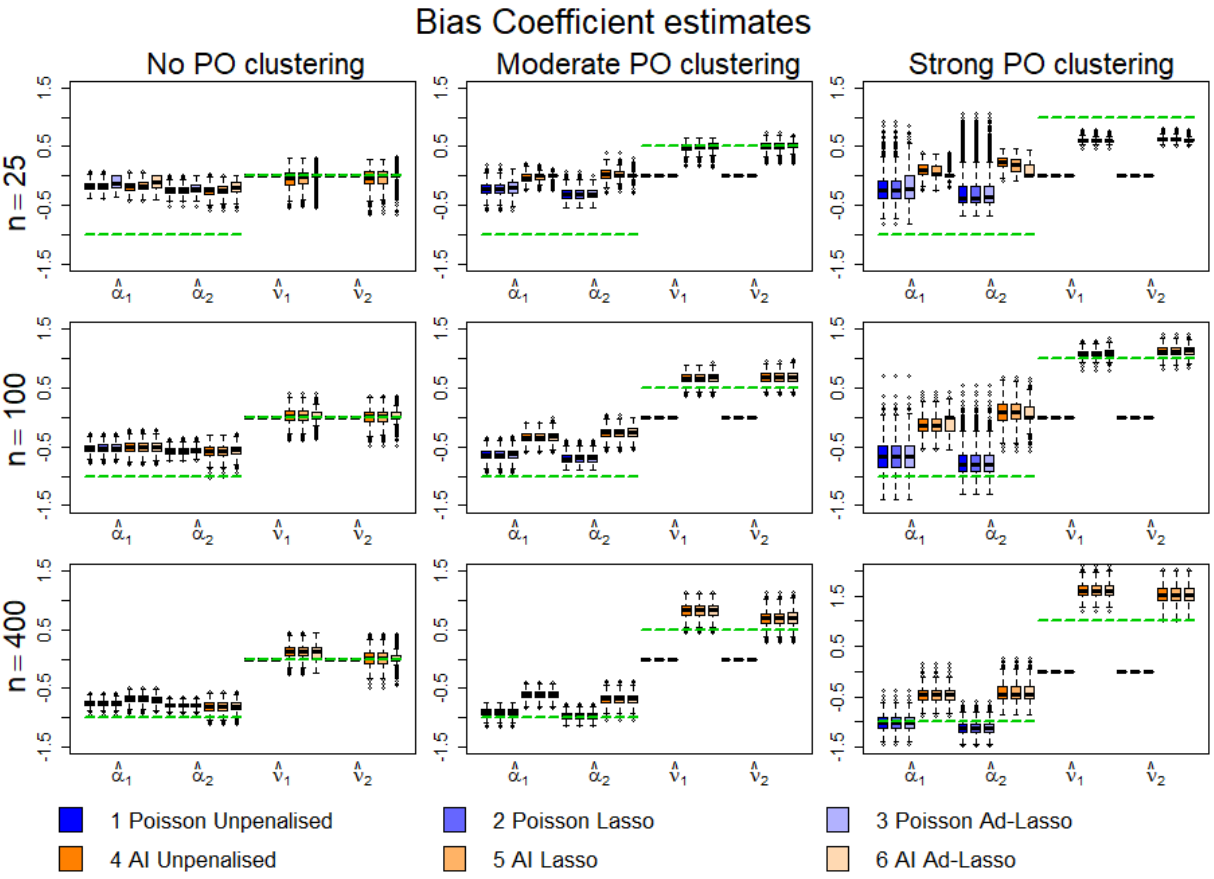


Figure A3: Boxpots of coefficient estimates corresponding to the two observer bias and two area-interaction covariates for the six models described in Table 1 of the manuscript. The green lines indicate the true values of the parameters used to generated the simulated data.

Although there could be improvements to be made by better disentangling the spatial signal from the environmental and bias signals, the incorporation of an area-interaction term when presence-only points are clustered beyond what can be explained by the other model covariates still represents a significant improvement over standard implementations of combined likelihood approaches to species distribution modelling.

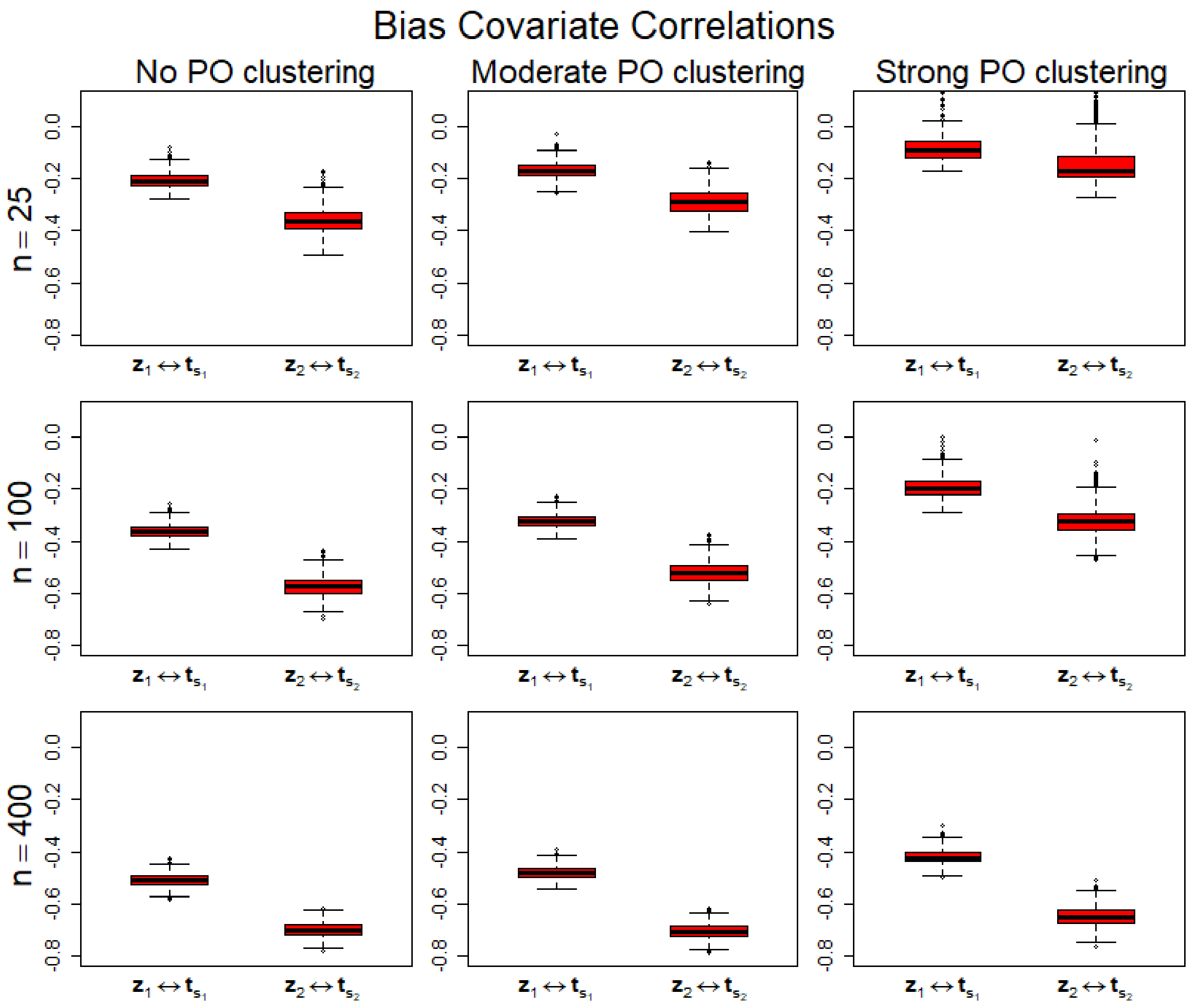


Figure A4: Boxpots of Pearson correlation coefficients between the observer bias and area-interaction covariates for both simulated data sources.
