## Supplementary material for "Combining multiple data sources in species distribution models while accounting for spatial dependence and overfitting with combined penalised likelihood maximisation": Optim Lasso demo

### Demonstration of Optim Lasso Functions

*25 June 2019*

This document demonstrates use of the `R` functions that were used to perform the simulations and the lynx analysis. These functions are contained in the file `Share Functions.R`. Here, we illustrate use of the functions through one of the simulations – in particular, the first simulation of presence-only patterns with 100 points that exhibit moderate clustering, as determined by the `strength` object. To ensure that roughly 100 points are generated, we set the intercepts for the two patterns with `int1` and `int2`. To ensure moderate clustering, we set the value of the interaction coefficient `intcoef`.

```
source("Share Functions.R")
seeduse = 1
strength = 3
N = 100
int1 = -2.25
int2 = -2.4
intcoef = 0.5
```

The functions we demonstrate make use of some functions of other packages, which we now load:

```
library(mvtnorm)
library(sp)
library(lattice)
library(spatstat)
library(data.table)
library(raster)
library(ROCR)
```

#### Generating environmental variables

First, we will generate the environmental covariates revelant to the simulated species distribution. First, we define our observation window as $[0, 30] \times [0, 30]$ and generate uniformly random "centers", from which we will randomly draw a subset to serve as the centres of habitat quality patches drawn from a truncated Gaussian distribution. These centers are set to be a distance of at least 1 unit from the window edge.

```
set.seed(3)
n_centers = 2000
centers = matrix(runif(n_centers, 1, 29), n_centers/2, 2)
plot(centers[,1], centers[,2], xlim = c(0, 30), ylim = c(0, 30),
     xlab = "X", ylab = "Y", main = "Population of Patch Centers",
     asp = 1)
```

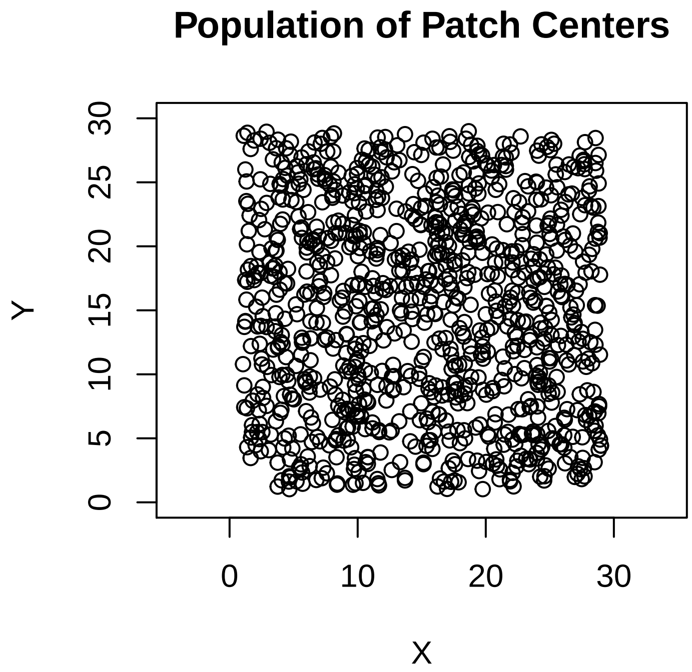


We also define a regular grid of quadrature points at a 0.5-unit spatial resolution throughout the observation window. It is at these quadrature points that we will measure the generated covariates.

```
quad = expand.grid(seq(0, 30, 0.5), seq(0, 30, 0.5))
names(quad) = c("X", "Y")
```

We now use the `makecovar` function to generate the two environmental covariates which define the species distribution:

```
set.seed(1)
x1 = makecovar(100)
set.seed(16)
x2 = makecovar(250)
```

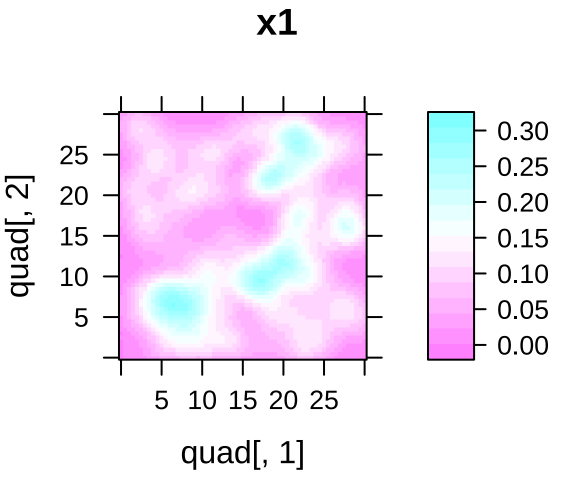


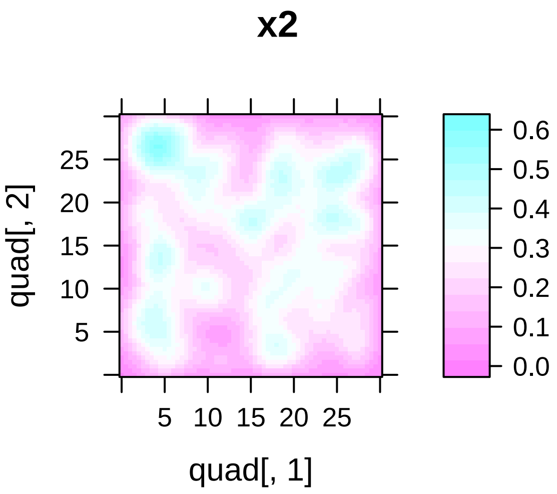


We now generate a covariate `catcov` to serve as a categorical covariate, the distance from which will be used to bias one of the presence-only data sources. First, we generate a continuous covariate `contcov` and categorise it through a threshold to generate `catcov`:

```
set.seed(18)
contcov = makecovar(100, centers, quad)
catcov = rep(0, length(contcov)) # categorical for PO bias
catcov[contcov >= quantile(contcov, 0.75)] = 1
levelplot(catcov ~ quad[,1] + quad[,2], main = "categorical covariate", asp = "iso")
```

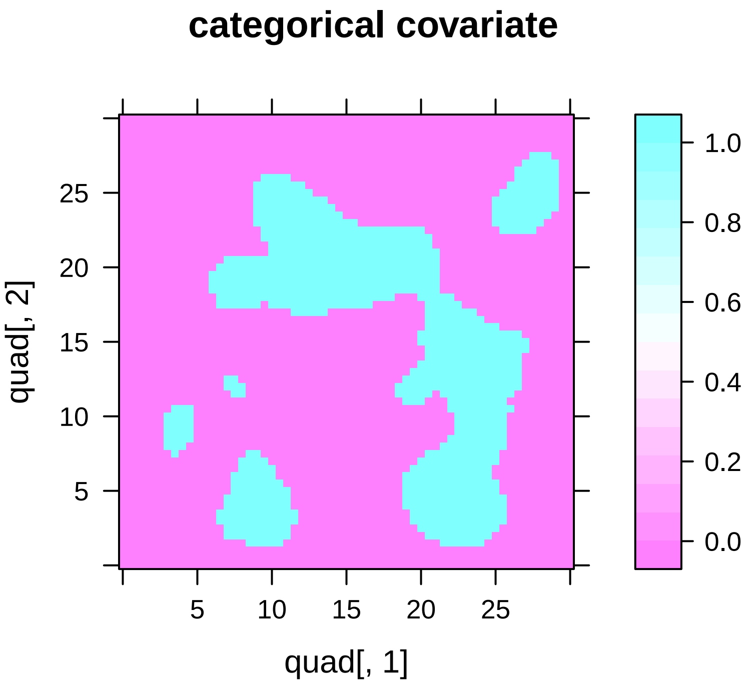


We will also generate a covariate that defines detection probability for the occupancy data:

```r
set.seed(20)
z3 = makecovar(120, centers, quad) # detection for occupancy
levelplot(z3 ~ quad[,1] + quad[,2], main = "z3: detection covariate", asp = "iso")
```

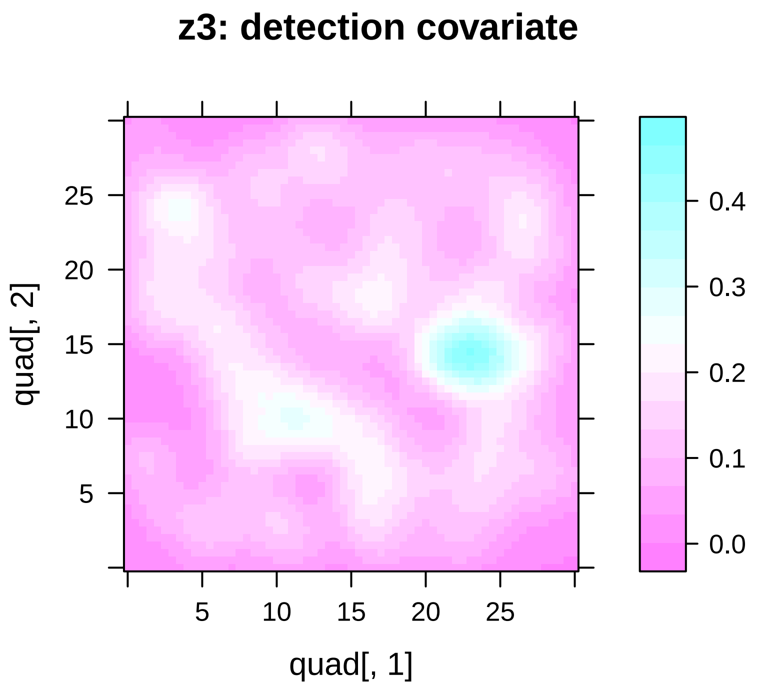


We will also generate two "dummy" covariates that we will include in models which do not impact the true species distribution. This allows us to check the performance improvement that results from applying lasso-type penalties in the combined penalised likelihood.

```r
set.seed(5)
d1 = makecovar(50)
set.seed(15)
d2 = makecovar(50)
```

The seeds were chosen for all of the above covariates to ensure that none of the covariates are highly correlated:

```r
covframe = data.frame(x1, x2, contcov, catcov, z3, d1, d2)
cor(covframe)
```

```
##                x1         x2   contcov     catcov           z3
## x1      1.0000000 0.33335245 0.2718674 0.16609826  0.290223677
## x2      0.3333524 1.00000000 0.3256248 0.18697137  0.395409641
## contcov 0.2718674 0.32562479 1.0000000 0.80270078  0.377083449
## catcov  0.1660983 0.18697137 0.8027008 1.00000000  0.307465718
## z3      0.2902237 0.39540964 0.3770834 0.30746572  1.000000000
## d1      0.1447537 0.02424448 0.1194169 0.07558192 -0.006555065
## d2      0.1955840 0.31041880 0.2102573 0.09821886  0.241281489
##                  d1         d2
## x1       0.144753684  0.19558399
## x2       0.024244476  0.31041880
## contcov  0.119416856  0.21025726
## catcov   0.075581920  0.09821886
```

```
## z3      -0.006555065  0.24128149
## d1       1.000000000 -0.13324797
## d2      -0.133247973  1.00000000
```

We will now simulate a road network in order to generate distances from the nearest road as a bias covariate for the first presence-only data source. The `addnetwork` function creates a network of roads. This function proceeds by adding $n$ small road segments in random directions (normally distributed with a standard deviation of 3 initially centered at a value set by the `angle` argument). It randomly splits the road into junctions (a *parent* road and a *child* road), with the probability of a split set by the `p_split` argument and the angle of the child road a minimum of `min_angle` from the previous road. The number of levels (*generations*) of road splits is set by `max_levels`, with subsequent splits for each generation having split probabilities multiplied by a value set by the `split_decay` argument (to allow different levels of the network to have more or less frequent splits) and the standard deviation of angles between road segments multiplied by a value set by the `sdmult` argument (to allow different levels of the network to have different amounts of "wiggliness").

Here, we will generate 3 road networks of 3 levels each and combine them. Road segments belonging to levels 1, 2, and 3 are plotted in black, red, and green, respectively. The `fixnodes` function ensures that junctions aren't too close – any junctions within a small distance of each other are combined, producing a more realistic road network.

```
set.seed(3)
network1 = addnetwork(0, 15, angle = 0, p_split = 0.03, split_decay = 0.2,
                      nmult = 0.6, min_angle = 60, sdmult = 1.5, max_levels = 3)
set.seed(12)
network2 = addnetwork(0, 15, angle = 40, p_split = 0.035, split_decay = 0.25,
                      nmult = 0.6, min_angle = 60, sdmult = 1.5, max_levels = 3)
set.seed(5)
network3 = addnetwork(0, 15, angle = -40, p_split = 0.015, split_decay = 0.2,
                      nmult = 0.6, min_angle = 60, sdmult = 1.5, max_levels = 3)
combined = c(network1, network2, network3)
combinednew = fixnodes(combined)
plotroadlist(combinednew, main = "Simulated Road Network", asp = 1)
```

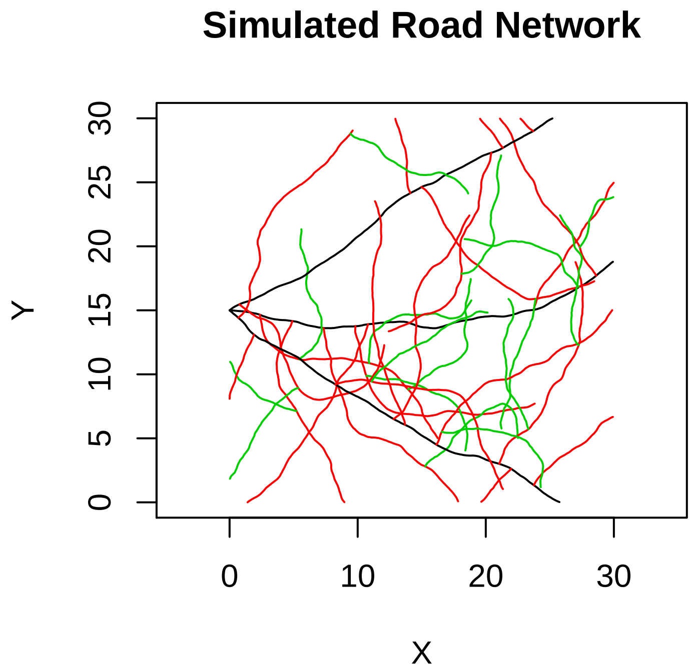


```
combinednewnodes = nodelist(combinednew)
```

We now generate distances to the nearest road in the simulated road network as a bias variable `d_rd` ($\mathbf{z}_1$ in the manuscript) for the first presence-only data source.

```
allpts = ppp(x = c(quad$X, combinednewnodes$X),
             y = c(quad$Y, combinednewnodes$Y),
             marks = c(rep("Quad", dim(quad)[1]), rep("Road", dim(combinednewnodes)[1])),
             window = owin(c(0, 30), c(0, 30)))
nndists = nndist(allpts, k = 1, by = as.factor(marks(allpts)))
d_rd = nndists[1:dim(quad)[1], 2] # distance to the nearest road
levelplot(d_rd ~ quad[,1] + quad[,2], main = "d_rd: bias for presence-only source 1", asp = "iso")
```

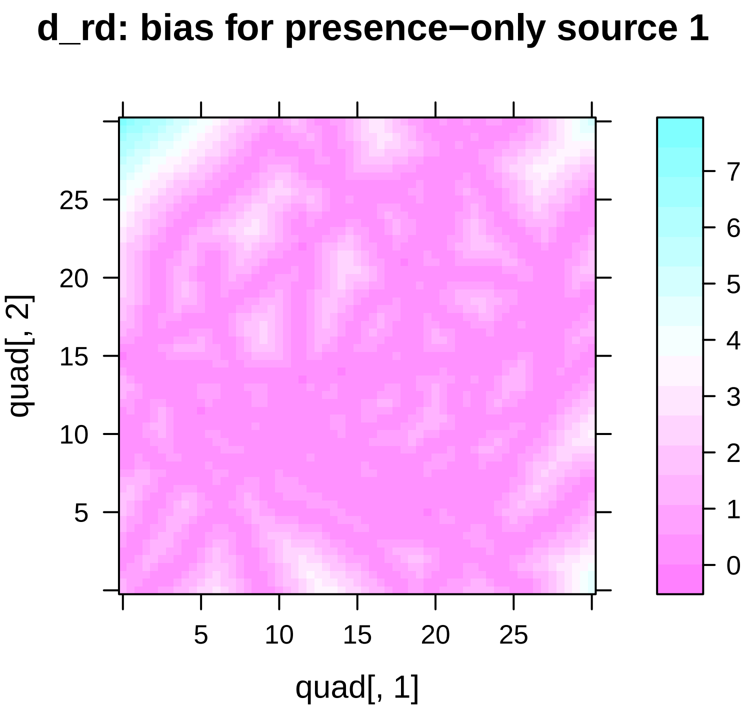


We also generate distance to the categorical variable `catcov` as a bias variable `d_cat` ($\mathbf{z}_2$ in the manuscript) for the second presence-only data source.

```
catpts = ppp(x = quad$X, y = quad$Y,
             marks = paste("Cat", catcov, sep = ""),
             window = owin(c(0, 30), c(0, 30)))
catdists = nndist(catpts, k = 1, by = as.factor(marks(catpts)))
d_cat = catdists[,2]
d_cat[catcov == 1] = 0 # distance to categorical variable
levelplot(d_cat ~ quad[,1] + quad[,2], main = "d_cat: bias for presence-only source 2", asp = "iso")
```

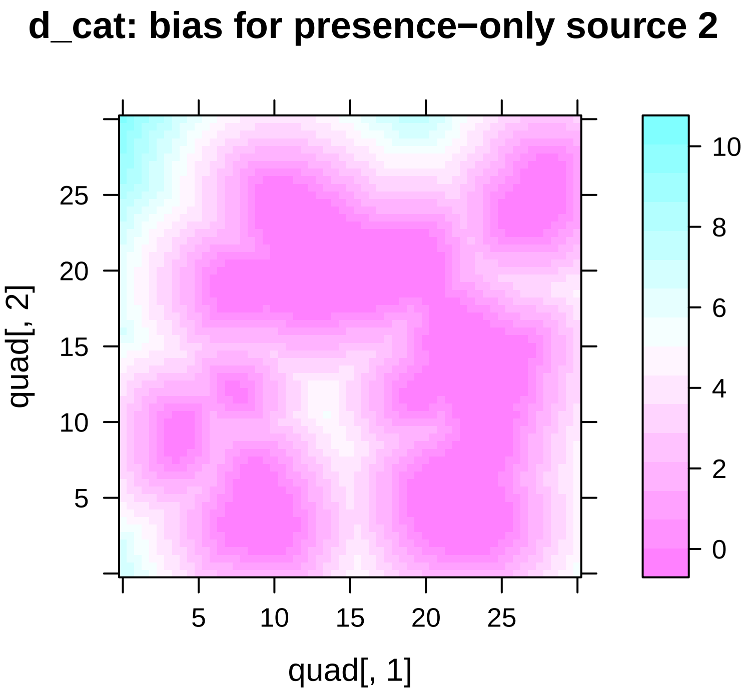


We now standardise the covariates to have mean 0 and variance 1, so that the lasso-type penalties operate fairly on the coefficients.

```
x1 = scale(x1)
x1sq = scale(x1^2)
x2 = scale(x2)
x2sq = scale(x2^2)
z3 = scale(z3)
rt_d_rd = scale(sqrt(d_rd))
rt_d_cat = scale(sqrt(d_cat))
d1 = scale(d1)
d1sq = scale(d1^2)
d2 = scale(d2)
d2sq = scale(d2^2)
```

#### Generating point patterns

We now generate point patterns for the true species as well as two presence-only data sources as a function of the generated environmental and bias covariates. We will also generate a site history for use in an occupancy model.

```
vmat = as.matrix(data.frame(1, x1, x1sq, x2, x2sq,
                            rt_d_rd, contcov, catcov, rt_d_cat))
env.mat = vmat[,1:5] # vmat without covariates for bias
```

The true species intensity `sp_int` will make use of the environmental variables and an intercept (contained in `env.mat`):

```
sp_coef = c(2.15, 0.5, -0.3, 0.5, -0.3)
sp_int = exp(env.mat %*% sp_coef)
levelplot(sp_int ~ quad$X + quad$Y, asp = "iso", main = "True intensity")
```

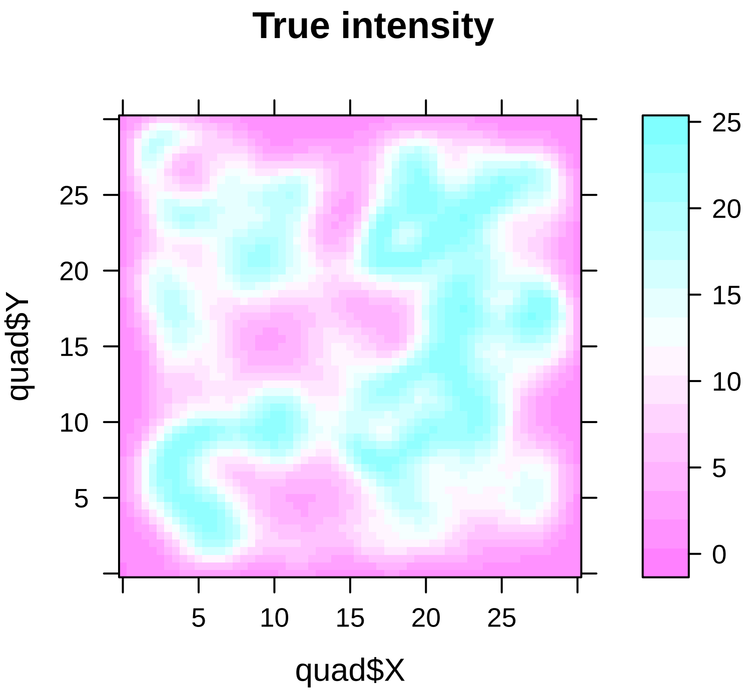


The two biased presence-only intensities will have the same environmental coefficients as the true intensity, but will be biased by `rt_d_rd` (pattern 1) and `rt_d_cat` (pattern 2):

```
po1_coef = c(int1, 0.5, -0.3, 0.5, -0.3, -1)
po1_int = exp(vmat[,1:6] %*% po1_coef)
levelplot(po1_int ~ quad$X + quad$Y, asp = "iso",
          main = "Presence-only source 1 intensity")
```

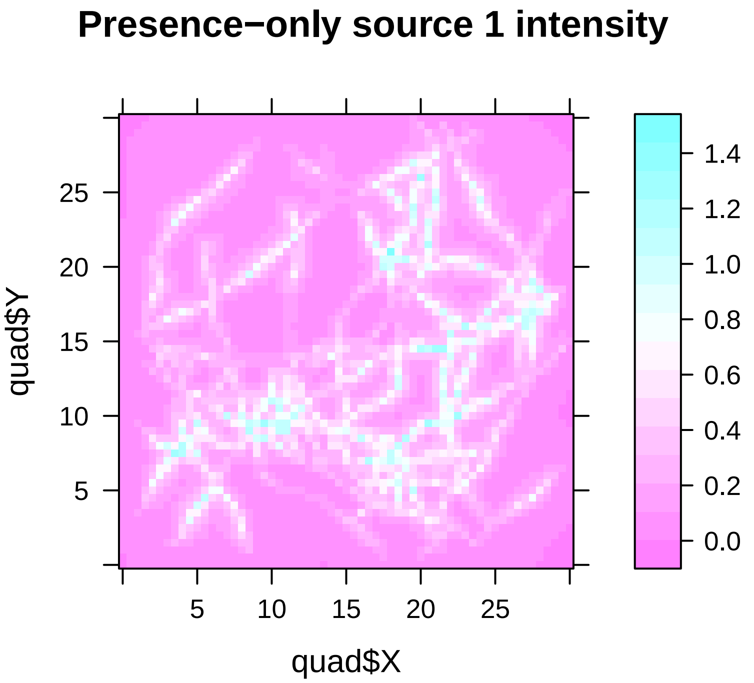


```r
po2_coef = c(int2, 0.5, -0.3, 0.5, -0.3, -1)
po2_int = exp(vmat[,c(1:5, 9)] %*% po2_coef)
levelplot(po2_int ~ quad$X + quad$Y, asp = "iso",
          main = "Presence-only source 2 intensity")
```

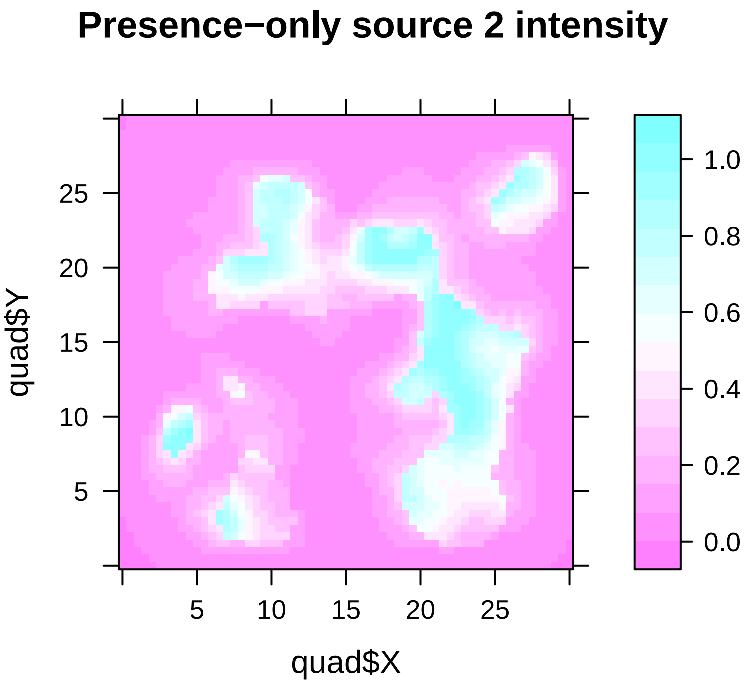

We can see that presence-only source 1 has a strong bias toward the road network, and presence-only source 2 has a strong bias toward the categorical covariate. The spatstat package requires us to format these intensity surfaces as images in order to generate point patterns:

```
sp_int_im = as.im(data.frame(x = quad$X, y = quad$Y, z = sp_int))
po1_int_im = as.im(data.frame(x = quad$X, y = quad$Y, z = po1_int))
po2_int_im = as.im(data.frame(x = quad$X, y = quad$Y, z = po2_int))
```

We now generate the true species pattern with the rpoispp function of spatstat.

```
set.seed(seeduse)
sp_sim = rpoispp(sp_int_im) # true species pattern
sp_sim$n
```

```
## [1] 10349
```

```
sp_xy = data.frame(X = sp_sim$x, Y = sp_sim$y)
plot(sp_sim)
```

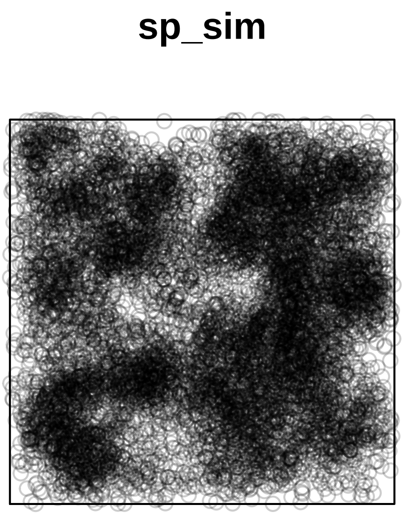


To fit models, we need to collect the values of the environmental and bias covariates at the grid of quadrature points and the species locations. First, we generate a data frame of quadrature point locations and associated covariates (quads), and then interpolate values at the species locations (sp_env) using the newenv.var function:

```
quads = data.frame(X = quad$X, Y = quad$Y, X1 = x1, X2 = x1sq,
                   X3 = x2, X4 = x2sq, Z3 = z3, rt_d_rd = rt_d_rd,
                   rt_d_cat = rt_d_cat, D1 = d1, D2 = d1sq,
                   D3 = d2, D4 = d2sq)
sp_env = newenv.var(sp.xy = sp_xy, env.grid = quads,
                    env.scale = 0.5, coord = c("X", "Y"), file.name = NA)
```

```
## Calculating species environmental data for variable: X1
```

```
## Calculating species environmental data for variable: X2
## Calculating species environmental data for variable: X3
## Calculating species environmental data for variable: X4
## Calculating species environmental data for variable: Z3
## Calculating species environmental data for variable: rt_d_rd
## Calculating species environmental data for variable: rt_d_cat
## Calculating species environmental data for variable: D1
## Calculating species environmental data for variable: D2
## Calculating species environmental data for variable: D3
## Calculating species environmental data for variable: D4
```

To generate the presence-only sources with clustering, we will proceed with an iterative algorithm that sequentially samples points from the species pattern `sp_sim` proportionally to conditional intensities and adds them to the set of observed locations. To compute conditional intensities, we essentially add an extra covariate – the *point interactions* of the pattern. For an area-interaction model, we can compute the point interaction at a location $s$ by drawing circles of a given radius $r$ around the location $s$ and compute the proportion of the circle that overlaps with the circles of radius $r$ around the other points in the pattern $\mathbf{s}$. This proportion is edge-corrected for points near the border of the observation window. By setting the coefficient for this interaction term to be positive, we increase the conditional intensity at points that have high point interactions (i.e. points that are clustered around other observed points), encouraging clustering.

First, we initialise the interaction covariate to be a vector of zeros, and we sample the first point from the pattern `sp_sim` at random, with sampling probabilities proportional to the computed intensities at the locations in `sp_sim`.

```
po1_X = data.frame(Intercept = 1, sp_env[,c(3:6, 8)], Interaction = 0)
po1_beta = c(int1, 0.5, -0.3, 0.5, -0.3, -1, intcoef)
po1_intensity = exp(as.matrix(po1_X) %*% po1_beta)
PO1_rows = c() #vector indicating the sampled rows
X_add = sample(1:sp_sim$n, 1, prob = po1_intensity)
PO1_rows = c(PO1_rows, X_add)
PO1 = sp_sim[PO1_rows]
```

We then compute the point interactions for all of the points in `sp_sim` based on the first sampled point, and standardise this vector as we standardised the environmental and bias covariates.

```
ints = evalInteraction(PO1, sp_sim, interaction = AreaInter(1), correction = "border")
stand_ints = scale(ints)
```

We now iterate through steps to add the remaining points. We first redefine the vector of point interactions, compute the conditional intensity `int_i` (setting it to 0 for the points already sampled to ensure they aren't resampled), sampling a new point based on probabilities proportional to the updated conditional intensities, and recomputing standardised point interactions based on the new set of points.

```
for (i in 2:N)
{
  po1_X$Interaction = stand_ints
  int_i = exp(as.matrix(po1_X) %*% po1_beta)
  int_i[PO1_rows] = 0
  X_add = sample(1:sp_sim$n, 1, prob = int_i)
  PO1_rows = c(PO1_rows, X_add)
  PO1 = sp_sim[PO1_rows]
  ints = evalInteraction(PO1, sp_sim, interaction = AreaInter(1), correction = "border")
  stand_ints = scale(ints)
}
po1_xy = data.frame(X = PO1$x, Y = PO1$y)
plot(PO1)
```

## PO1

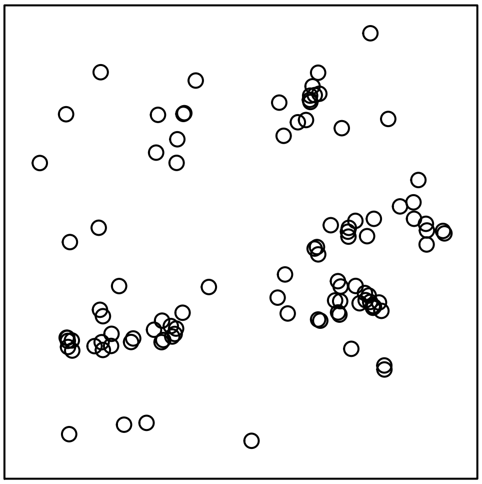

We follow the same procedure to sample points for the second presence-only data set, biased by distance from the categorical covariate.

```
po2_X = data.frame(Intercept = 1, sp_env[,c(3:6, 9)], Interaction = 0)
po2_beta = c(int2, 0.5, -0.3, 0.5, -0.3, -1, intcoef)
po2_intensity = exp(as.matrix(po2_X) %*% po2_beta)
PO2_rows = c()
X_add = sample(1:sp_sim$n, 1, prob = po2_intensity)
PO2_rows = c(PO2_rows, X_add)
PO2 = sp_sim[PO2_rows]
ints = evalInteraction(PO2, sp_sim, interaction = AreaInter(1), correction = "border")
stand_ints = scale(ints)
for (i in 2:N)
{
  po2_X$Interaction = stand_ints
  int_i = exp(as.matrix(po2_X) %*% po2_beta)
  int_i[PO2_rows] = 0
  X_add = sample(1:sp_sim$n, 1, prob = int_i)
  PO2_rows = c(PO2_rows, X_add)
  PO2 = sp_sim[PO2_rows]
  ints = evalInteraction(PO2, sp_sim, interaction = AreaInter(1), correction = "border")
  stand_ints = scale(ints)
}
po2_xy = data.frame(X = PO2$x, Y = PO2$y)
plot(PO2)
```

## PO2

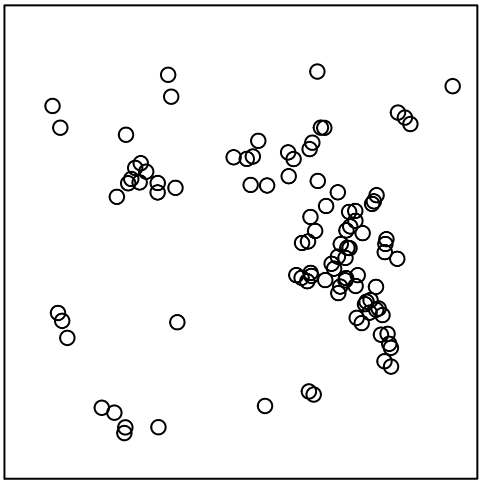

We now set up a grid of sites to be used in an occupancy model. We set up 100 sites along a regular $1.5 \times 1.5$-unit grid:

```
occ_sites = expand.grid(seq(1.5, 28.5, 3), seq(1.5, 28.5, 3))
occ_x = occ_sites[,1]
occ_y = occ_sites[,2]
```

We then determine whether each site is occupied by the species or not by computing distances between the site centers and the nearest point in the true species pattern. Any site center within a distance of 0.18 units from the nearest point in `sp_sim` is considered to be occupied, and the other sites are considered unoccupied.

```
sp_and_occ = ppp(x = c(occ_x, sp_sim$x), y = c(occ_y, sp_sim$y),
                 marks = c(rep("Occ", length(occ_x)), rep("Sp", sp_sim$n)),
                 window = owin(c(-0.25, 30.25), c(-0.25, 30.25)))
dist_sp_occ = nndist(sp_and_occ, k = 1, by = as.factor(marks(sp_and_occ)))
# compute distance to nearest species
sp_occ_dists = dist_sp_occ[1:length(occ_x), 2]
# species considered present at site if nearest one is within 0.25 units
occ_present = as.numeric(sp_occ_dists <= 0.18)
```

To determine detection probability at the sites, we map them to the quadrature points, which contain the values of all of the covariates. We use the inverse of the complementary log-log function to define detection probability. We set detection probability to 0 for those sites considered unoccupied by the species.

```
occ_paste = paste(occ_x, occ_y)
quad_paste = paste(quad$X, quad$Y)
occ_quadrow = match(occ_paste, quad_paste)
p_detect = cloglog inv(z3[occ_quadrow])*occ_present
```

We now generate a history `sim_history` of detections and non-detections at each site for 5 visits, randomly generating detections and non-detections according to the computed detection probabilities:

```
n_visits = 5
sim_history = matrix(as.integer(matrix(rep(p_detect, times = n_visits),
                                       length(occ_x), n_visits) >
                                  matrix(runif(length(occ_x)*n_visits),
                                         length(occ_x), n_visits)),
                     length(occ_x), n_visits)
site_sum = apply(sim_history, 1, sum)
plot_xy = expand.grid(seq(0, 30, 0.5), seq(0, 30, 0.5))
plot_x = plot_xy[,1]
plot_y = plot_xy[,2]
plot_id = paste(plot_x, plot_y)
occ_id = paste(occ_x, occ_y)
occ_match = match(occ_id, plot_id)
plot_z = rep(NA, length(plot_x))
plot_z[occ_match] = site_sum
levelplot(plot_z ~ plot_x + plot_y, asp = "iso", cuts = 5,
          col.regions = rainbow(6, start = 0.03, end = 0.17)[6:1])
```

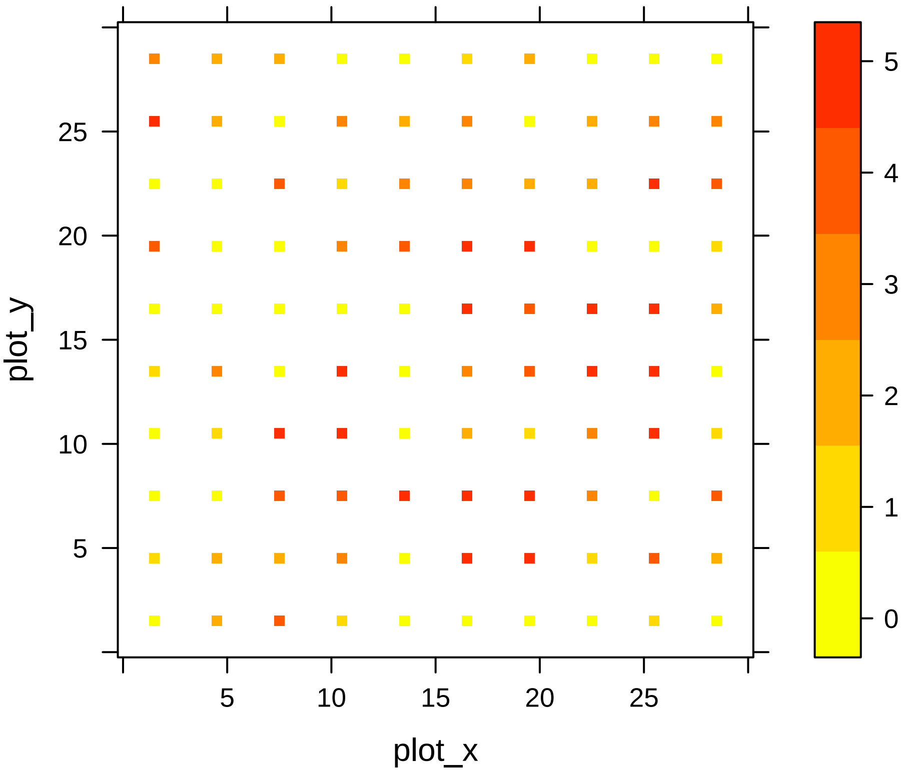


#### Fitting models

To fit models, we need the values of the environmental and bias variables at the species locations in the two presence-only data sets as well as at the locations of the occupancy sites. Since the presence-only locations are subsets of the pattern `sp_sim`, for which we already derived the values of the covariates in `sp_env`, we just need to extract the appropriate rows of the `sp_env` data frame.

```
po1_env = sp_env[PO1_rows,]
po2_env = sp_env[PO2_rows,]
```

Finally, we generate the values of the covariates at the occupancy sites by subsetting the appropriate rows from `quads`, since the occupancy sites were set up to be directly on one of the quadrature points:

```
occ_env = quads[occ_quadrow,]
```

We now create some objects that will be used in the multiple calls to the `comb_lasso` function to fit models. In particular, we specify the environmental formula used to parameterise $\boldsymbol{\beta}$ as well as the formulas for the various bias terms used to parametrise $\boldsymbol{\alpha}$. We specify the interaction radius for models fitted with the area-interaction likelihood with the `int.radius` object.

```
env_formula = ~ X1 + X2 + X3 + X4 +
  D1 + D2 + D3 + D4
bias_formula = list(~ rt_d_rd, ~ rt_d_cat, ~ Z3)
n.fits = 1000
int.radius = c(1, 1)
tol = 1.e-9
b.min = 1.e-6
link = "cloglog"
sp_res = 0.5
```

We first generate models with the inhomogeneous Poisson point process model presence-only likelihood. The arguments are as follows:

- `intercept_env`: whether to include (1) or not (NA) an intercept for the environmental component of the model
- `intercept_bias`: a list of indicators for inclusion of intercepts in the various bias components of the model
- `quad_data`: a data frame for the quadrature data with columns containing values for all covariates
- `sp_data`: a list of data frames for the species data with columns containing values for all covariates
- `sp_y`: a list of objects containing the response variables for each species data source. For presence-only data sources, this is a vector of 1s. For occupancy models, it is a matrix of detections and non-detections.
- `dat.type`: a vector of species data types, with "PO" indicating presence-only and "Occ" indicating occupancy.
- `coord`: names for the columns that include the $x$- and $y$-coordinates in the `sp_data` objects
- `sp_res`: spatial resolution to be used in fitting the presence-only point process models
- `penalty_vec`: a pre-supplied vector of penalties to be used. If `NULL`, this vector will be computed automatically
- `alpha`: the elastic net coefficient. If `alpha` is set to 1, a lasso penalty will be applied. If `alpha` is set to 0, a ridge regression penalty will be applied. If `alpha` is between 0 and 1, an elastic net penalty will be applied, with the value of `alpha` controlling the lasso portion of the penalty.
- `gamma`: exponent for adaptive lasso weights
- `init.coef`: initial coefficients used to determine adaptive lasso weights. If `NULL`, a standard lasso is fitted
- `standardise`: a logical value indicating whether the covariates should be standardised to have mean 0 and variance 1 prior to model fitting
- `criterion`: which criterion is used to determine the optimal model among the fitted models. Set to "BIC" by default but AIC ("AIC"), corrected AIC ("AICc") and the Hannah-Quinn criterion ("HQC") are also permitted.
- `family`: the canonical link family used in the point process models – set to Poisson by default
- `tol`: a tolerance threshold controlling convergence of the lasso algorithm. The lasso is fitted by iteratively reweighted least squares, and if the difference in likelihoods between successive iterations falls below `tol`, the model is considered to have converged
- `b.min`: the smallest absolute value of $\beta$ permitted. The function sets any estimated coefficient with an

absolute value less than `b.min` to 0

- `max.it`: the maximum number of iterations of the iteratively reweighted least squares algorithm for the lasso
- `n.fits`: the number of models to fit, if `penalty_vec` is not specified.
- `noshrink`: a vector indicating the covariates not to be penalised. If `NULL`, only the intercept terms will be unpenalised.
- `method`: the optimisation method used by `optim`
- `link`: the link function for the detection covariate – either "cloglog" or "logit" is permitted.
- `site.area`: the area of the occupancy sites
- `area.int`: a logical value indicating the form of the presence-only likelihood components. If `TRUE`, area-interaction model likelihoods are used for each presence-only source. If `FALSE`, an inhomogeneous Poisson PPM likelihood is used instead.
- `r`: The radius of the point interactions if `area.int` is set to `TRUE`
- `wt.vec`: A vector of source weights to be applied to each likelihood component. By default, set to `NULL` such that the weight is 1 for each source.
- `pen.min`: The smallest non-zero penalty to be applied. By default, this is set to 0.01.

```r
comb_ppm = comb_lasso(env_formula = env_formula,
                      bias_formula = bias_formula,
                      intercept_env = 1,
                      intercept_bias = list(NA, NA, NA),
                      quad_data = quads,
                      sp_data = list(po1_env, po2_env, occ_env),
                      sp_y = list(rep(1, nrow(po1_env)), rep(1, nrow(po2_env)), sim_history),
                      dat.type = c("PO", "PO", "Occ"),
                      coord = c("X", "Y"),
                      sp_res = sp_res,
                      penalty_vec = NULL,
                      alpha = 1, gamma = 0, init.coef = NA, standardise = TRUE,
                      family = "poisson", tol = tol, b.min = b.min,
                      max.it = 25, n.fits = n.fits, noshrink = NULL, method = "BFGS",
                      link = link, site.area = pi*0.18^2,
                      area.int = FALSE, r = NULL)
```

```
## [1] "Output saved in the file Quad0.5.RData"
```

To fit a model with an adaptive lasso penalty, we specify a value for the adaptive weight exponent `gamma` as well as a vector of initial coefficients `init.coef`:

```r
comb_ad_ppm = comb_lasso(env_formula = env_formula,
                      bias_formula = bias_formula,
                      intercept_env = 1,
                      intercept_bias = list(NA, NA, NA),
                      quad_data = quads,
                      sp_data = list(po1_env, po2_env, occ_env),
                      sp_y = list(rep(1, nrow(po1_env)), rep(1, nrow(po2_env)), sim_history),
                      dat.type = c("PO", "PO", "Occ"),
                      coord = c("X", "Y"),
                      sp_res = sp_res,
                      penalty_vec = NULL,
                      alpha = 1, gamma = 1, init.coef = comb_ppm$betas[,1],
                      standardise = TRUE,
                      family = "poisson", tol = tol, b.min = b.min,
                      max.it = 25, n.fits = n.fits, noshrink = NULL, method = "BFGS",
                      link = link, site.area = pi*0.18^2,
                      area.int = FALSE, r = NULL)
```

```
## [1] "Output saved in the file Quad0.5.RData"
```

We can fit models with an area-interaction term by setting the `area.int` argument to `TRUE` and supplying a vector of radii for the point interactions to the `r` argument:

```
comb_ai = comb_lasso(env_formula = env_formula,
                      bias_formula = bias_formula,
                      intercept_env = 1,
                      intercept_bias = list(NA, NA, NA),
                      quad_data = quads,
                      sp_data = list(po1_env, po2_env, occ_env),
                      sp_y = list(rep(1, nrow(po1_env)), rep(1, nrow(po2_env)), sim_history),
                      dat.type = c("PO", "PO", "Occ"),
                      coord = c("X", "Y"),
                      sp_res = sp_res,
                      penalty_vec = NULL,
                      alpha = 1, gamma = 0, init.coef = NA, standardise = TRUE,
                      family = "poisson", tol = tol, b.min = b.min,
                      max.it = 25, n.fits = n.fits, noshrink = NULL, method = "BFGS",
                      link = link, site.area = pi*0.18^2,
                      area.int = TRUE, r = c(int.radius, NA))
```

```
## [1] "Output saved in the file Quad0.5.RData"
## Calculating point interactions
## Calculating point interactions
```

We can also fit models using just a single source of data, though such models are not considered in the manuscript. As an example, the following fits models using only the first presence-only data source:

```
po1_ppm = comb_lasso(env_formula = env_formula,
                         bias_formula = list(bias_formula[[1]]),
                         intercept_env = 1,
                         intercept_bias = NULL,
                         quad_data = quads,
                         sp_data = list(po1_env),
                         sp_y = list(rep(1, nrow(po1_env))),
                         dat.type = c("PO"),
                         coord = c("X", "Y"),
                         sp_res = sp_res,
                         penalty_vec = NULL,
                         alpha = 1, gamma = 0, init.coef = NA, standardise = TRUE,
                         family = "poisson", tol = tol, b.min = b.min,
                         max.it = 25, n.fits = n.fits, noshrink = NULL, method = "BFGS",
                         link = link, site.area = pi*0.18^2,
                         area.int = FALSE, r = NULL)
```

```
## [1] "Output saved in the file Quad0.5.RData"
```

#### Exploring Fitted Models

The created objects contain a lot of information about the fitted models, including the fitted coefficients for all models (`betas`), the vector of penalties (`penalty_vec`), the penalised likelihoods (`pen_likelihoods`), the values of the various criteria to choose the lasso penalty (`criterion_matrix`), the fitted intensity (`mu`), occupancy probability (`psi`), and detection probability (`p_detect`) of the optimal model, the runtime (`runtime`) and the data used in the model fitting.

We can examine the fitted coefficients for the model which optimised BIC:

```
comb_ppm$beta
```

```
##   Intercept          X1          X2          X3          X4          D1
## -2.74433124  0.83598140 -0.40490977  0.75378210 -0.41387000 -0.06719399
##          D2          D3          D4     rt_d_rd    rt_d_cat          Z3
## -0.08224648 -0.02960134 -0.01281307 -0.53227289 -0.72330739  0.93211371
```

```
comb_ad_ppm$beta
```

```
##   Intercept          X1          X2          X3          X4          D1
## -2.73712493  0.83219212 -0.40722838  0.73666386 -0.39834125 -0.03346626
##          D2          D3          D4     rt_d_rd    rt_d_cat          Z3
## -0.08568419  0.00000000  0.00000000 -0.53113523 -0.72172138  0.92847091
```

```
comb_ai$beta
```

```
##   Intercept          X1          X2          X3          X4          D1
## -2.69681660  0.41282015 -0.20131157  0.43632514 -0.23054465  0.00000000
##          D2          D3          D4     rt_d_rd Interaction    rt_d_cat
## -0.07917936 -0.08556838  0.02062521 -0.26909853  0.67793812 -0.23966321
## Interaction          Z3
##  0.69291701  0.92757279
```

```
po1_ppm$beta
```

```
##    Intercept           X1           X2           X3           X4
## -4.186028204  1.812580070 -0.925689009  1.386875530 -0.755216546
##           D1           D2           D3           D4      rt_d_rd
##  0.148990250 -0.398124979 -0.153440610 -0.004871374 -0.663559637
```

We may wish to see some details of the *regularisation path* of fitted models. To view a plot of the coefficients as the penalty increases, we can use the `plotpath` function. For the combined model with IPPPM likelihoods fitted with an adaptive lasso penalty:

```
plotpath(comb_ad_ppm)
```

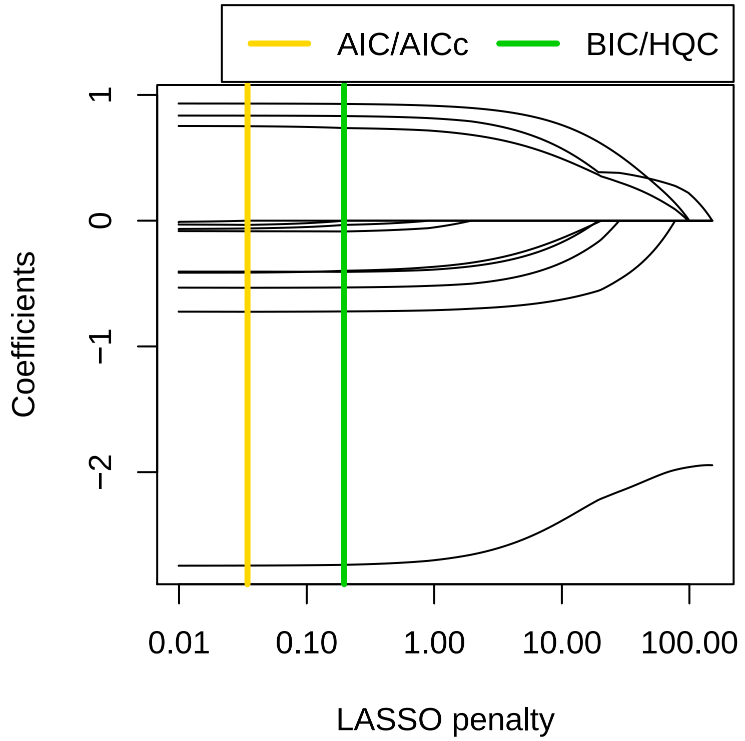


We can wish to view the regularisation path without the intercept in this case, as it is not peanlised. To do so, we supply the index of the intercept to the `v.cut` argument:

```
plotpath(comb_ad_ppm, v.cut = which(names(comb_ppm$beta) == "Intercept"))
```

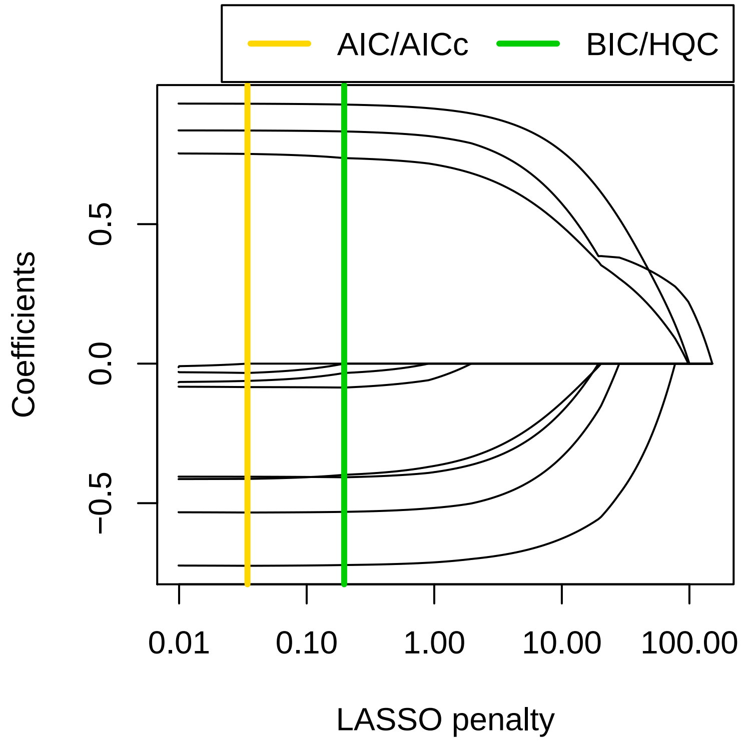

We can also plot the value of the criterion used to choose the best model (in this case BIC) for all fitted models with the `criterion_curve` function:

```
criterion_curve(comb_ad_ppm)
```

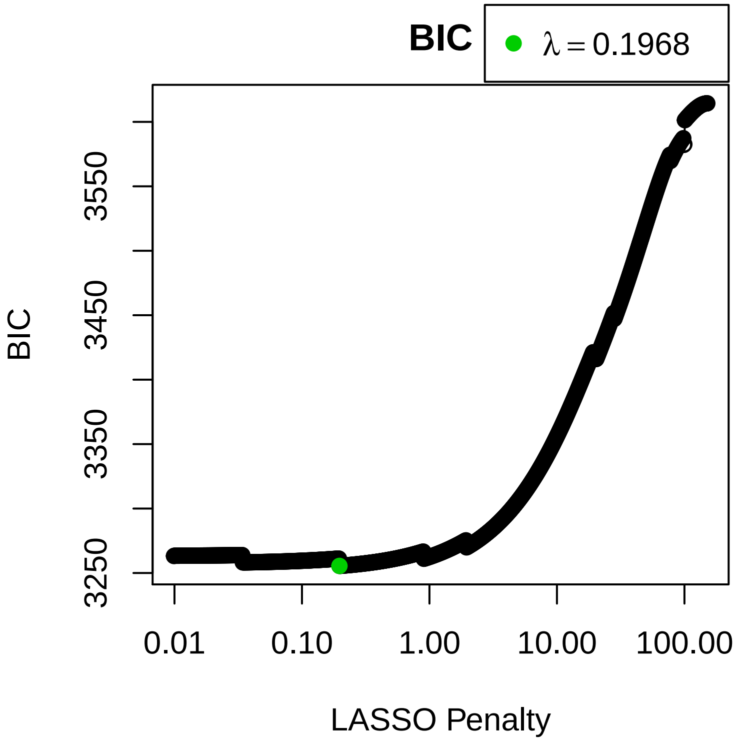


The `plotfit` function allows us to produce heatmaps for various quantities for our optimal fitted model. To plot the fitted intensity for the optimal model:

```
plotfit(comb_ad_ppm)
```

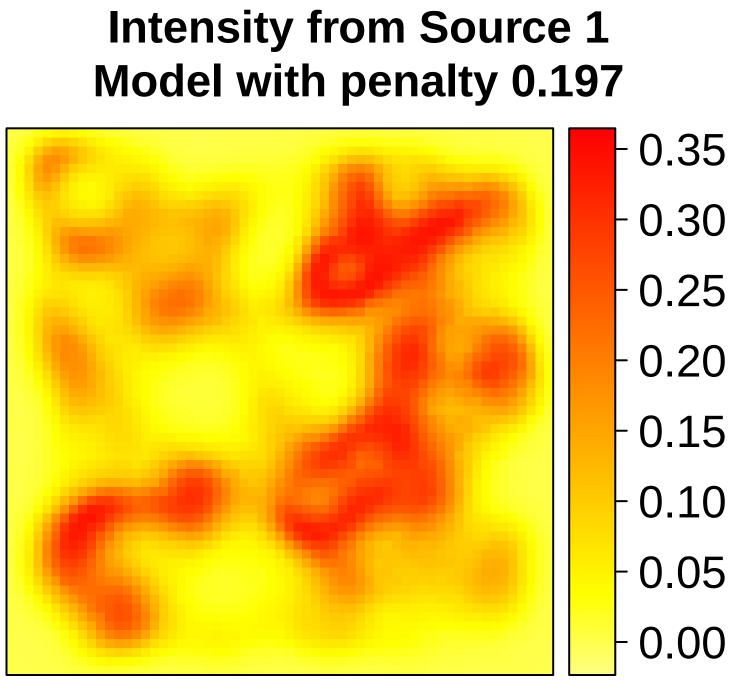


To plot the occupancy probability:

```
plotfit(comb_ad_ppm, z = "occupancy", link = "cloglog")
```

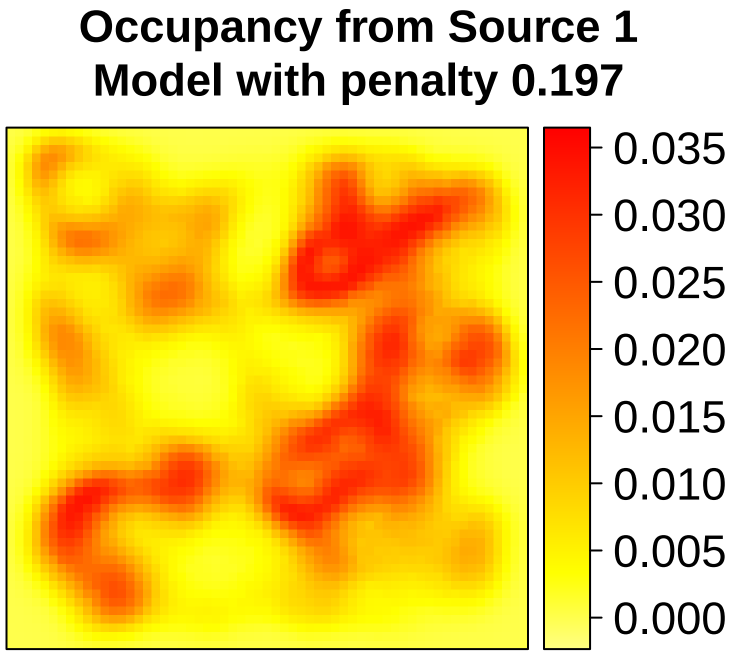


To plot the estimated bias for each source:

```
plotfit(comb_ad_ppm, z = "bias", source = 1)
```

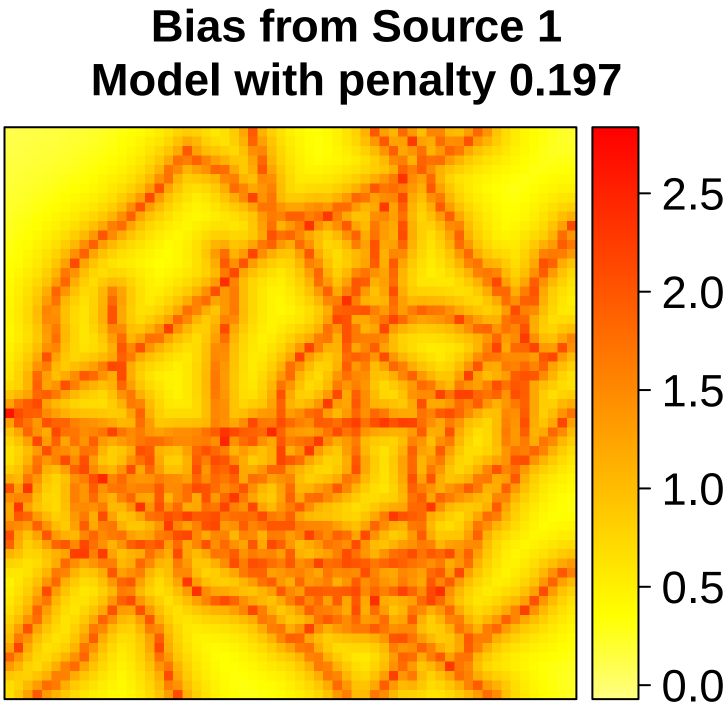


```
plotfit(comb_ad_ppm, z = "bias", source = 2)
```

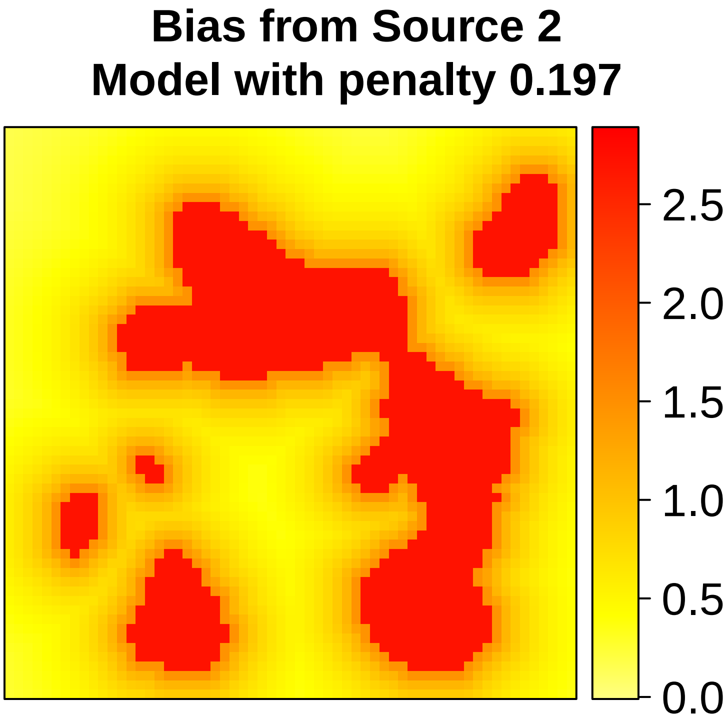

We can also choose to plot these quantities for any of the fitted models. For example, the fitted intensity for the 800th fitted model:

```
plotfit(comb_ad_ppm, model = 800)
```

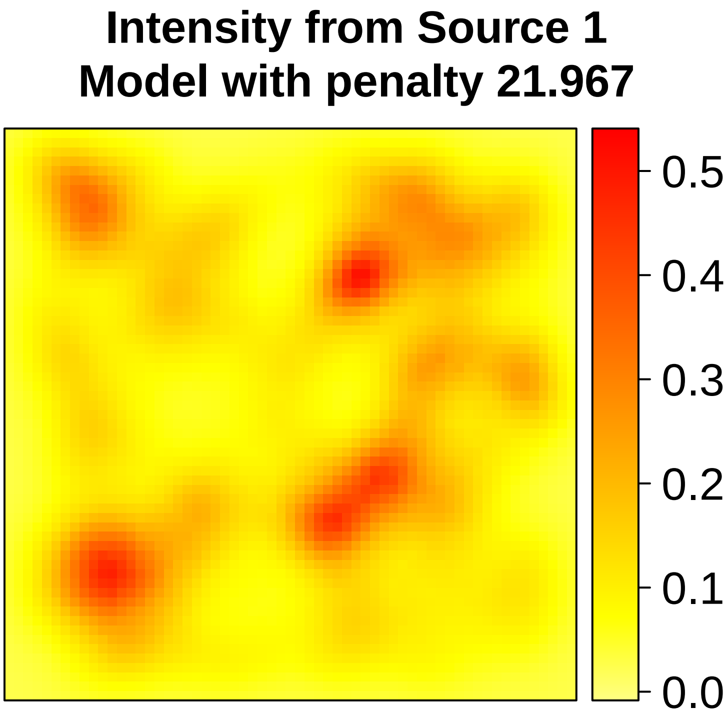
